# Genome-wide Cell-free DNA Fragmentation Profiling at Transcription Start Sites Detects Cancer Without Cancer-Type-Specific Region Selection

**DOI:** 10.64898/2026.06.22.733692

**Authors:** Bram Pronk, Stavros Makrodimitris, Saskia M Wilting, Marcel J T Reinders

## Abstract

Accurate discrimination between healthy individuals and patients with cancer using minimally invasive liquid biopsies could improve cancer diagnosis and monitoring. Circulating cell-free DNA (cfDNA) is a promising biomarker, since its fragmentation patterns reflect chromatin organization at regulatory regions such as transcription start sites (TSSs). Classification approaches typically rely on hypothesis-driven selection of genomic regions based on literature or external tissue data. They thereby assume that tumor-derived cfDNA carries the diagnostic signal, potentially overlooking a systemic, genome-wide shift in the cfDNA pool. We present a data-driven framework that selects TSSs directly from cfDNA sequencing data, with every modeling choice made within nested cross-validation. For breast and colorectal cancer, it reached an AUROC of 0.966, against 0.891 for tumor-specific chromatin accessibility regions, 0.666 for gene-expression regions and 0.673 for aneuploidy-based ichorCNA. Across seven cancer types it reached 0.969, and 0.95 for cancer types left out of training. On an additional hepatocellular carcinoma dataset, the AUROC was 0.74, while both hypothesis-driven baselines performed at chance. The model’s discriminatory signal is spread over thousands of TSSs and may reflect the cellular origin of cfDNA. These results support data-driven locus selection as an alternative to cancer-type-specific region sets for cfDNA-based cancer detection.

## 1. Introduction

Liquid biopsy-related biomarkers based on circulating cell-free DNA (cfDNA) have demonstrated substantial potential in precision oncology. They provide minimally invasive access to tumor-associated molecular information, including tumor burden, subtype, and therapy resistance [1, 2, 3]. Genome-wide sequencing of cfDNA has shown that fragmentation patterns are informative of nucleosome organization [4], DNA methylation [5], tissue of origin [6], and copy number variation [7]. All of these have been linked to underlying physiology [8] as well as the genomic landscape of tumors in patients with cancer [2, 9].

Cell-free DNA fragmentation patterns reflect nucleosome occupancy in the originating cells [4, 10], providing a genome-wide readout of chromatin organization across contributing cell populations [8, 11]. Recent studies have leveraged this principleby interrogating cfDNA fragmentation profiles in both genome-wide settings [3] and at transcriptionally relevant genomic loci. These loci include Transcription Start Sites (TSSs) [4], Transcription Factor Binding Sites (TFBSs) [12, 2] and disease-specific regions-of-interest (ROIs) such as DNase I hypersensitive sites and enhancer regions [13]. In these approaches, coverage or fragmentomic features are aggregated over predefined regions, either large genomic bins [3] or sets of ROIs. Differences between patients with cancer and healthy controls are then quantified to enable classification. Such strategies have yielded biological insight into nucleosome organization in cancer and have demonstrated discriminatory potential in specific disease settings [14].

To decide which ROIs to investigate, such methods are typically hypothesis-driven: genomic regions are selected a priori based on knowledge about the disease profile and the affected cell type. This selection uses external datasets such as tumor-specific chromatin accessibility profiles, DNA methylation maps, or differential gene expression data. However, several factors can negatively influence the efficacy of these approaches. First, defining appropriate ROIs requires prior knowledge of the cancer type and for an understudied cancer type often requires additional large-scale molecular profiling, restricting clinical transferability. Second, even with an appropriate set of cancer-specific regions, fragmentation signals at selected loci may be attenuated in samples with low circulating tumor DNA (ctDNA) fractions, as encountered in early-stage disease or minimal residual disease settings, or in low-pass sequencing settings. Third, focusing on tumor- or tissue-derived regions imposes constraints on where in the genome cancer-associated cfDNA changes must appear. However, these constraints might be too strict. Recent work has shown that much of the increased cfDNA in patients with cancer does not come from the tumor itself, reflecting systemic shifts in cell turnover [15]. Fragmentomic alterations previously linked to cancer are also present in inflammatory conditions independent of malignancy [16]. Consequently, cancer-associated fragmentation signals may reflect distributed alterations in cellular turnover and chromatin organization across multiple contributing cell types, in addition to circulating tumor DNA itself.

These considerations led us to hypothesize that a data-driven approach would better capture the potentially distributed nature of the cancer-associated fragmentation signal. Such an approach identifies discriminative loci directly from cfDNA data rather than relying on predefined cancer- or tissue-specific region sets. By leaving locus selection unconstrained, it avoids prior assumptions about which genomic regions carry diagnostic information. It can also accommodate signals arising from systemic alterations beyond the tumor itself. In this study, we select informative TSSs directly from cfDNA sequencing data, with every modeling choice made within a nested cross-validation. We show that this strategy reaches a higher area under the receiver operating characteristic curve (AUROC) than established hypothesis-driven ROI selection methods derived from gene expression and chromatin accessibility data, as well as copy number–based approaches. This holds both for breast and colorectal cancer and across seven cancer types. We also show that it largely retains its performance on cancer types left out of training. The signal it relies on is spread over thousands of TSSs and may reflect shifts in the cellular origin of cfDNA rather than tumor-specific regions. This suggests that integrating weak signals across the genome can make a priori region selection unnecessary.

## 2. Methods

### 2.1. Datasets

We used two publicly available Whole-Genome Sequencing (WGS) datasets of plasma cfDNA. Anonymized fragment-level data from both cohorts were obtained from FinaleDB [17] and converted to BAM format using a modified implementation of the Fragmentstein framework [18]. All samples were processed against the hg38 reference genome. GC bias correction was performed using GCfix [19] with default parameters.

The primary cohort is from Cristiano et al. [3]. From the 534 records retrieved, we kept one sequencing record per participant. We excluded non-plasma samples, repeated runs, lung cancer samples that were not treatment-naïve, and one duodenalcancer case. Additionally, we excluded 30 samples from healthy individuals that were sequenced much more deeply than the rest of the cohort (Supplementary Figure S1, Supplementary Table S1). The 214 healthy participants from Cristiano et al. were then split into two groups. The first 50 in the order of retrieval from FinaleDB were held out as a reference set. They were used exclusively for filtering low-quality TSSs (Section 2.2) and as a panel of normals for ichorCNA (Section 2.5). Thus the final modeling cohort comprised the other 164 healthy participants and 203 treatment-naïve patients with cancer spanning seven cancer types (mean coverage: ~2.48x, range: 1.06x - 5.25x).

As an additional dataset, we also used the cohort described by Jiang et al. [20]. It consists of 90 patients with hepatocellular carcinoma (HCC) and 135 non-HCC participants: 32 healthy subjects, 67 patients with chronic hepatitis B and 36 with hepatitis B-associated cirrhosis (mean coverage: ~2.11x, range: 1.20x - 4.84x). Both cohorts were sequenced on the Illumina HiSeq 2000. However, they differ in read length (100 bp paired-end for Cristiano et al., 75 bp for Jiang et al.) and in extraction and library preparation protocols (Supplementary Table S2). The fragment-length distributions and other per-sample measures of the two cohorts are compared in Supplementary Figure S2a, b.

The data of both cohorts were collected with informed consent and ethical approval in the original studies, which are described in [3] and [20]. This study is a secondary analysis of the anonymized fragment-level data released through FinaleDB [17]; no participants were recruited and no identifiable information was accessed.

### 2.2. ROI Feature Extraction

#### 2.2.1. TSS Windows

To capture a disease-related signal in a data-driven way, we required a predefined search space of genomic regions. This space had to be biologically meaningful, structured enough to permit feature extraction at individual loci, and not selected for any cancer type or tissue. Transcription Start Sites (TSSs) meet these criteria (Figure 1). Active TSSs exhibit a well-characterized chromatin architecture consisting of a nucleosome-depleted region (NDR) flanked by positioned nucleosomes. This configuration is reflected in cfDNA coverage profiles [4, 21, 2].

**Fig. 1:**
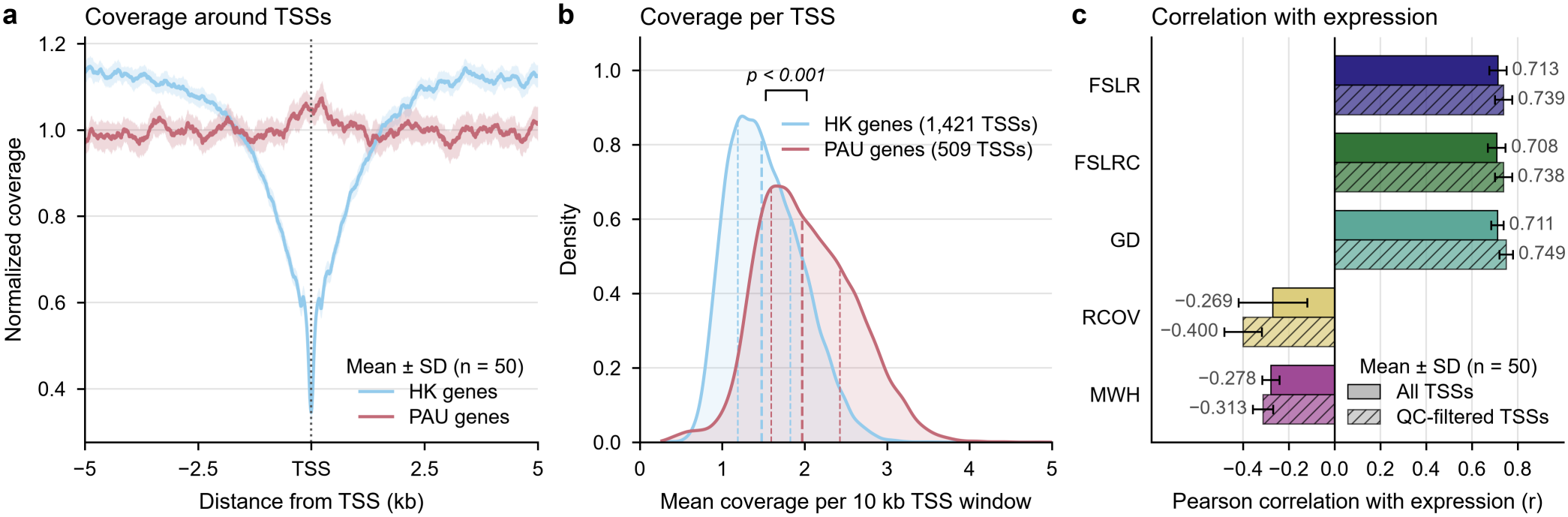
cfDNA fragmentomic features at TSSs and their correlation with gene expression in the panel of normals. a) Normalized cfDNA coverage around the TSSs of housekeeping (HK, blue; 1,421 TSSs) and unexpressed (PAU, red; 509 TSSs) genes, counting the TSSs that pass the quality control. For each sample, the coverage summed over the TSSs of a gene set is divided by its mean over the 10 kb window, so the mean is 1. Lines show the mean over the 50 samples; shaded bands show *±* 1 standard deviation. b) Distribution of the mean coverage in the 10 kb window of each HK and PAU TSS, pooled over the 50 samples; dashed lines mark the quartiles. c) Pearsoncorrelation of each feature with gene expression in a peripheral blood mononuclear cell (PBMC) reference (log_2_ TPM per gene group, Esfahani et al.), computed per sample and averaged over the 50 samples; error bars show *±* 1 standard deviation. Solid bars use all TSSs; hatched bars use only the TSSs that pass the quality filter. RCOV and MWH correlate negatively with expression by construction. *Alt text:* Graphs labelled a to c on cell-free DNA around transcription start sites. a, coverage dips sharply at the start sites of highly expressed housekeeping genes and stays flat at those of unexpressed genes. b, coverage across the window is lower for the housekeeping genes. c, bar chart in which three fragmentation features correlate positively and two negatively with gene expression.

To extract features from TSSs, we obtained TSS coordinates from Ensembl Biomart v112 (GRCh38.p14) for 18,534 protein-coding genes located on autosomes. For genes with multiple annotated TSSs, we used the one labeled as “canonical”. In the case of multiple “canonical” TSSs, we kept the one with the lowest genomic coordinate. Fragmentomic features were calculated within a 10k bp region of interest (ROI) centered around each TSS, thus covering the 5 kb up- and downstream of the TSS [2].

#### 2.2.2. Quality control

Several quality control steps were applied to filter out problematic ROIs. Regions overlapping the ENCODE blacklist [22] or additional low-mappability and artefactual regions defined by GCparagon [23] were excluded (n = 1,210; Supplementary Figure S3a, b). The 3,286 TSS windows with another TSS within 5 kb were excluded, because their coverage pattern can be influenced by the chromatin state of the nearby TSS. Finally, we computed the standard deviation of coverage of each 10 kb TSS ROI across the 50 held-out healthy samples of the Cristiano et al. cohort. To ensure robustness under low-pass sequencing, we excluded the 0.5% of ROIs with the highest standard deviation (n = 70 that no other filter had removed; Supplementary Figure S3c). This removes only extreme outliers. In total, 4,308 of the 18,534 windows were removed (258 by more than one filter), and 14,226 TSS ROIs remained. Supplementary Figure S3 shows the same filters on all 19,376 protein-coding TSSs, including the 842 on chromosome X.

#### 2.2.3. Fragmentomic features

For each retained ROI, five fragmentomic features were extracted (Supplementary Table S3). The Fragment Short/Long Ratio is defined by either absolute fragment counts (FSLR) or coverage (FSLRC) [14, 3]. The other three are the Griffin Difference (GD) [14], relative nucleosome-depleted region coverage (RCOV) [24] and the Max Wave Height (MWH) [14]. RCOV and MWH are by their definitions not calculated over the entire 10k bp window. For the MWH, the promoter region was defined as −120 bp to +30 bp relative to the TSS. The first nucleosome in the gene body was defined as +31 bp to +195 bp [14]. All five features were computed relative to the coverage or fragment counts within their own window (Supplementary Table S3). Each sample’s feature vector was then z-scored across its TSSs to adjust for the overall coverage level of each sample.

To illustrate the expected cfDNA coverage patterns at TSSs,we used two reference gene sets. The first set comprises 2,176 housekeeping (HK) genes consistently expressed across 52 tissues and cell types [25]. The second comprises 751 non-tissue-specific genes with very low protein expression (nTPM *<* 2.5) in the Human Protein Atlas [26], termed Protein Atlas Unexpressed (PAU) genes [23].

#### 2.2.4. Correlation with gene expression

To test whether the features reflect gene expression, we correlated them with expression levels in peripheral blood mononuclear cells (PBMCs), following Esfahani et al. [27]. Genes were ordered based on their mean log(TPM) expression in PBMCs and summarized in groups of ten consecutive genes [27]. For each sample of the panel of normals, we averaged each feature over the TSSs of the genes in a group. The Pearson correlation between these group means and the group expression was then computed. The correlations were averaged over the 50 samples. This was repeated after the quality control above, keeping only the groups in which at most one gene was removed and using the remaining TSSs.

### 2.3. Data-driven ROI Selection

We implemented a data-driven feature selection framework that does not rely on predefined, hypothesis-driven region sets. It treats each TSS as an independent candidate biomarker, to identify a subset of genomic regions that optimally distinguish healthy cfDNA samples from those of patients with cancer. For each sample and each TSS, the five fragmentomic features (FSLR, FSLRC, GD, MWH, RCOV) were computed. This yielded five feature matrices *X*_*p*_ ∈ ℝ^*S×T*^, where *S* denotes the number of samples and *T* =14,226 the number of TSSs. For each feature, ROIs whose values were identical to those of another ROI in all samples were removed. This left 14,221 ROIs for GD, the feature used by the final models, and between 14,132 and 14,220 for the other features. We ranked these candidate regions in each *X*_*p*_ by their ability to separate healthy from cancer samples, using training samples only (Section 2.6). For this we implemented four ranking functions: (1) the p-value of a two-sided Mann-Whitney U (MWU) test, (2) the absolute difference between the group means, and(3) the absolute log_2_ ratio of the group means. Function (4) is a weighted score, the product of (3) and the − log_10_(*p*) of the MWU test. Random selection of *k* TSSs served as a control to the ranking functions.

Functions (3) and (4) are heuristic rankings, not fold changes in the usual sense: the features are z-scored within each sample (Section 2.2), so a group mean can be negative or close to zero. A ratio of group means with opposite signs has no logarithm, and such TSSs receive a score of zero (about 5% of TSSs). A mean close to zero makes the ratio large even when the difference between the groups is small.

The top *k* TSSs were selected to construct a feature vector that was used in a classifier. Here, *k* was evaluated over a range spanning 10 to all 14,221 retained TSSs. The ranking function, the fragmentomic feature, and *k* were treated as tunable hyperparameters optimized using a cross-validation framework (Section 2.6, Supplementary Table S4).

### 2.4. Comparison to hypothesis-driven approaches

The data-driven framework was benchmarked against two hypothesis-driven region selection strategies based on RNA sequencing (RNA-seq) and assay for transposase-accessible chromatin using sequencing (ATAC-seq) data.

#### 2.4.1. RNA-seq baseline

For the RNA-seq–based approach, we used the 792 blood-specific nucleosome-depleted regions defined by Zhu et al. [24]. These regions correspond to promoters and/or exon-intron junctions of 265 genes highly expressed in whole blood and lowly expressed across 20 solid tumor types. Expression was determined using data from GTEx [28] and TCGA [29], respectively. For each region, we only compute the relative nucleosome-depleted region coverage (RCOV) feature as defined by the authors in [24]. This feature was chosen because its definition takes the specific region type (promoter versus intron–exon junction) into account.

#### 2.4.2. ATAC-seq baseline

For the ATAC-seq based approach, we used the transcription factors (TFs) identified by Ulz et al. [12]. These TFs have binding sites that are differentially accessible in colorectal cancer (CRC) or breast cancer (BRCA) samples relative to whole blood (411 TFs for CRC and 447 for BRCA). Their union, *n* = 504, was used for classification. The provided genomic coordinates of the top 1000 binding sites (based on ChiP-seq signal [30]) per transcription factor (TF) were converted to hg38 coordinates using UCSC liftover. Following Ulz et al., we constructed a composite coverage profile for each TF by aggregating cfDNA coverage across its 1000 binding sites. We computed one transcription factor accessibility (TFA) feature per profile (Supplementary Table S3). As the TFA feature was designed by Ulz et al. for these profiles, we used only this feature, as for the RNA-seq baseline. As for the TSSs (Section 2.3), regions whose values were identical to those of another region in all samples were removed before classification, which left 781 of the 792 RNA-seq regions and 497 of the 504 ATAC-seq features.

#### 2.4.3. Number of regions and sets of TSSs

To test whether the fixed number of regions puts the baselines at a disadvantage, we also ran both with the number of regions *k* as an additional hyperparameter. This hyperparameter was optimized in the inner loop (Section 2.6). The regions were ranked by the four ranking functions of Section 2.3, and *k* ranged from 10 to all 781 (RNA-seq) or 497 (ATAC-seq) regions.

We also tested whether aggregating the TSSs into predefined sets improves on single TSSs. The 14,221 TSSs were grouped into the 50 MSigDB Hallmark gene sets [31] (3,431 TSSs in total) and into the 39 autosomal chromosome arms. Each of the five features was computed per set, on the pooled coverage profiles or fragment counts or as the average of the per-TSS values. These set-level features went through the same classification procedure as the baselines, on the same outer folds (Methods 2.6). Each feature was also computed on all 14,221 TSSs pooled, which gives one value per sample. This value served as the score directly. Its sign was set on the training samples of each outer fold, so that higher scores indicate cancer.

### 2.5. Comparison to ichorCNA

We additionally compared the ROI-based models to genome-wide somatic copy number calling and tumor fraction estimation using ichorCNA [7]. Analyses were conducted using default parameters with two modifications. First, a panel-of-normals was constructed using the 50 held-out healthy samples from the Cristiano et al. cohort, which were sequenced similarly to the patients, to reduce noise and batch effects. Second, we followed the approach from Moldovan et al. [32] and lowered the maximum copy number to 3 to improve robustness under low tumor fraction conditions. For each sample, the tumor fraction corresponding to the solution with the highest log-likelihood was reported. For the comparisons to ichorCNA, the estimated tumor fraction was used as the classification score, with the threshold for 95% specificity derived from the training samples of each outer fold. The same setup was used to run ichorCNA on the samples of the Jiang et al. dataset.

### 2.6. Model training and evaluation

#### 2.6.1. Nested cross-validation

We evaluated the data-driven TSS selection framework for binary classification of cancer versus healthy cfDNA. We benchmarked it against hypothesis-driven region sets (Section 2.4) and copy number–based tumor fraction estimation via ichorCNA (Section 2.5). Unless stated otherwise, all supervised models were trained and evaluated using the same stratified nested cross-validation to prevent information leakage between model selection and performance estimation. The outer loop (10-fold) provided held-out test sets for estimating generalization performance. In the inner loop (3-fold), we optimized the hyperparameters for six classifiers: Support Vector Machine, Logistic Regression, Random Forest, K-Nearest Neighbors, a Multi-Layer Perceptron, and XGBoost (Supplementary Table S5). For the data-driven approach, the inner loop additionally performed a grid search to simultaneously select the optimal fragmentomic feature, the ranking function, and number of selected TSSs *k* (Supplementary Table S4). For the two hypothesis-driven baselines, the region set and feature were fixed, and the classifier and its hyperparameters were tuned. In an additional analysis, their number of regions *k* was tuned as well (Section 2.4). After inner-loop optimization, the model configuration with the highest mean inner-loop area under the receiver operating characteristic curve (AUROC) was retrained on the full outer training set. It was then evaluated once on the held-out outer test fold.

We report the generalization performance as the mean and standard deviation of the AUROC across the 10 outer folds. We also report the sensitivity at the 95% specificity cut-off. In each outer fold, this cut-off was the 0.95 quantile of the scores of the healthy training samples in the inner-loop predictions of the selected configuration. It was applied unchanged to the test fold. Each participant has one out-of-fold prediction, from the model of the outer fold in which the participant was held out. ROC and precision-recall curves of these pooled out-of-fold predictions are shown in Supplementary Figure S4. For ichorCNA, we ran the software as described in Section 2.5. For comparability, we report the mean and standard deviation of the AUROC over the same outer test folds.

The primary benchmark distinguished healthy controls (*n* = 164) from patients with breast or colorectal cancer (BRCA, n = 54; CRC, n = 26). We restricted the cancer class to these two malignancies because the hypothesis-driven region sets were originally derived for them. To evaluate the cancer-type-agnostic potential of the data-driven framework, we then ran a separate nested cross-validation against the healthy controls. In this task, the cancer class comprised all seven cancer types (PANCAN) in the Cristiano et al. cohort (n = 203). All splits were stratified by cancer type, so every type was present in training. A separate analysis leaves out one cancer type at a time (see below).

#### 2.6.2. Additional dataset

To evaluate generalization across cohorts, we applied the data-driven model to the Jiang et al. cohort (non-HCC participants n = 135; hepatocellular carcinoma n = 90). Hepatocellular carcinoma is a cancer type absent from the training data. The hyperparameter configuration for the data-driven method and the baselines was selected by 10-fold cross-validation on the full Cristiano et al. cohort. A single model was then retrained on the entire cohort using the selected configuration. That model was then applied, without further tuning, to the Jiang et al. samples. Its 95% specificity threshold was fixed on the Cristiano et al. cohort, from the cross-validated scores of the selected configuration. The AUROC was computed against all non-HCC participants and also per comparison group within the non-HCC participants (32 healthy subjects, 67 patients with chronic hepatitis B and 36 with hepatitis B-associated cirrhosis).

#### 2.6.3. Corrections and training within the Jiang et al. dataset

As we noticed systematic differences between the cohorts, we tried two corrections. First, the features of the Jiang et al. samples were standardized with the mean and standard deviation of all225 Jiang et al. samples instead of those of the Cristiano et al. cohort. This uses no labels. Second, five sequencing-derived measures were added to the TSS features (Section 2.7). Finally, to test whether the Jiang et al. samples contain the signal at all, we ran the complete nested cross-validation within the cohort alone. Every choice of the model (feature, ranking function, number of TSSs and classifier) was therefore made and trained on Jiang et al. samples only, and nothing of the Cristiano et al. model was reused. To see how the number of TSSs affects generalization, the model selection on the Cristiano et al. cohort and the nested cross-validation within the Jiang et al. dataset were also repeated at every fixed *k* of the grid (Figure 4c).

#### 2.6.4. Leaving out cancer types

Application to the Jiang et al. dataset confounds two sources of performance loss: an unseen cancer type and inter-cohort technical variability. To isolate cross-cancer generalization within a single technical setting, we performed a leave-one-cancer-type-out experiment using only the Cristiano et al. cohort. Model training and hyperparameter optimization were done using 10-fold cross-validation on the six other types and 130 healthy donors. The best model configuration was retrained on all these samples and applied once to all patients of the left-out type and the 34 remaining healthy donors, which were the same for every type. The AUROC of the same patients and donors in the nested pan-cancer cross-validation, where the type is in training, serves as reference. To test whether TSSs that score well in every training type generalize better, the configuration selected in all seven searches (GD, elasticnet logistic regression) was retrained with only the best-scoring 100, 1,000 or 5,000 TSSs. The TSSs were scored on the training samples by the absolute difference between group means, either over all training cancer types together or within each of the six training types, with the ranks averaged (Figure 4e).

#### 2.6.5. Statistical comparisons

All comparisons between methods were carried out on exactly the same outer fold splits; none used averaged ROC curves across folds. Within the Cristiano et al. cohort, the AUROC of the data-driven model was compared with that of each baseline with the corrected resampled *t*-test on the ten paired fold-level AUROCs. This test accounts for the overlapping training sets of the folds and gives the 95% confidence interval of the difference [33]; *p*-values were Holm-adjusted over the three baselines per task. On the Jiang et al. dataset, each method was applied as a single trained model. Here, the AUROCs were compared with the DeLong test for correlated ROC curves, which gives the confidence interval of the difference, with Holm adjustment over the two hypothesis-driven baselines. The same test compared the methods on each left-out cancer type, Holm-adjusted over the seven types. Their mean AUROC over the types was compared with a paired participant-level bootstrap. Confidence intervals (95%) of a single AUROC were obtained by a participant-level bootstrap (2,000 resamples). Cancer and non-cancer participants were resampled with replacement within their class. Confidence intervals (95%) of the sensitivity and specificity at the 95% specificity threshold were exact binomial (Clopper-Pearson) intervals of the number of correctly classified participants. Per cancer stage (I to IV) and within the healthy donors, the predictions of the data-driven model and of each baseline were compared per participant (one-sided Wilcoxon signed-rank test). These 15 comparisons per task were Holm-adjusted as one family. The comparisons per ichorCNA tumor-fraction group in Supplementary Figure S7c, f are exploratory and not adjusted.

### 2.7. Per-sample measures and covariate analyses

We collected eight per-sample measures for the Cristiano et al. cohort. Five are technical and demographic covariates (mean coverage, total fragment count, plasma volume, age and sex). Three are cancer-associated measures (mean fragment length, mean fragment GC content and cfDNA concentration). Age, plasma volume and cfDNA concentration were taken from Supplementary Table S1 of Cristiano et al. [3]. The other five were computed from the sequencing data with our pipeline. Sex was inferred from the reads on the sex chromosomes [34, 35], which agreed with the sex recorded by Cristiano et al. for every participant.

#### 2.7.1. Measures alone and classifiers on them

Each measure was evaluated alone, with its direction taken from the training samples of each outer fold. Classifiers were also trained as for the data-driven model (Section 2.6) on the five covariates, on the five sequencing-derived measures, and on all eight measures. The sequencing-derived measures can be computed in any cohort (coverage, fragment count, fragment length, GC content and sex). Plasma volume and age are not reported per participant for the Jiang et al. dataset. For it, a classifier on the sequencing-derived measures was trained on the pan-cancer task and applied once. The same was done for each of these measures alone and we also tested a combination with the data-driven TSS model, in which we concatenate the selected TSSs to the five sequencing-derived features.

#### 2.7.2. Relation to the data-driven model

We related the data-driven model to these measures in four ways:(i) The five covariates were added as features next to the selected TSSs, and the nested cross-validation was repeated, once with *k* selected in the inner loop and once for each fixed *k* of its grid. (ii) For each TSS, the GD value, z-scored within each sample, was regressed (ordinary least squares) on class alone and on class plus coverage, plasma volume, age and sex. The *t*-statistic of class was compared between the two to test whether the ranking of the TSSs was sensitive to the covariates. Fragment count was left out of this model as it was almost perfectly correlated with coverage. Missing values for age and plasma volume were set to the median. (iii) Within each class, the out-of-fold score of the model was correlated with each measure. (iv) Within each class and outer fold, the out-of-fold scores of the data-driven model were compared to those of each classifier on the measures. In these analyses, we used Spearman’s correlation and corrected p-values in (ii) and (iii) with the Benjamini-Hochberg procedure.

### 2.8. Sequencing depth and training-cohort size

To assess the robustness of the data-driven model to lower sequencing depths, every sample was downsampled uniformly to 1x and to 0.1x coverage. The features were extracted again, and the complete nested cross-validation was repeated at each depth on the same outer folds. The models trained at full depth were also applied unchanged, including their 95% specificity threshold, to the downsampled samples of the corresponding outer test folds. The paired AUROCs of the outer folds of every two depth arms (ten pairs per task) were compared with a two-sided Wilcoxon signed-rank test, with Holm adjustment over the ten pairs.

To test how the approach depends on the size of the training cohort, the complete nested cross-validation was repeated for both tasks in the Cristiano et al. cohort (BRCA/CRC and pan-cancer versus healthy) on stratified subsets. The subsets ranged from 90% down to 20%, in steps of 10%, and each was contained in the next larger one. The subsets of 20% to 40% were drawn three more times with other random seeds to assess variability in the lowest sample regimes. Each subset was compared with the full cohort on its own participants, who have an out-of-fold score from both procedures. The comparison used a participant-level bootstrap of the differencein pooled out-of-fold AUROC (5,000 resamples, stratified by class), with Holm adjustment over the subsets of a task.

### 2.9. Biological interpretation of the TSS ranking

#### 2.9.1. Stability of the ranking

For interpretation, we ranked the 14,221 TSSs of each task by Cohen’s *d*: the difference in mean z-scored GD between patients with cancer and healthy donors, divided by the pooled standard deviation. A TSS thus ranks highly only when this difference is large relative to the spread between samples. This ranking serves interpretation only. The models select their TSSs with the ranking functions of Section 2.3.

The stability of the ranking was measured as the Spearman correlation of Cohen’s *d* between the training partitions of the outer folds and between two halves of the cohort that share no samples (repeated 50 times). The final ranking was also compared to a ranking calculated with permuted labels (20 permutations). The stability of the highest-ranked TSSs was measured as the Jaccard index of the top-*k* sets (Supplementary Figure S5a, b). The absolute coefficients of the models refitted in each outer fold were compared between folds in the same way (Supplementary Figure S5c).

#### 2.9.2. Agreement across cancer types

To compare the TSS ranking between cancer types, the healthy donors and the patients of every cancer type were randomly split into halves (200 splits). Each comparison thus used two disjoint halves of the healthy donors, so that a shared healthy group could not make the rankings agree. Cohen’s *d* was computed for each half of a type against one half of the healthy donors. Spearman correlations were computed between halves of two types (Supplementary Figure S5d) and between each type and the six other types pooled (Section 3.7), with participant-level bootstrap 95% intervals (400 replicates). The comparison between pairs of types was repeated for the coefficients of the pan-cancer model configuration refitted on each half (20 splits; Supplementary Figure S5e). Correlations were divided by the geometric mean of the two within-type correlations (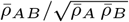 for types *A* and *B*, with means over the splits), so that a value of 1 means that two types agree as well as two halves of the same type.

#### 2.9.3. Gene sets

The ranked genes were tested for the overrepresentation of the 50 Hallmark sets [31], 965 Reactome pathways [36], and 738 C8 cell-type signatures from MSigDB [37] (version 2024.1), using the 14,221 TSSs as background. The 265 blood-specific promoter genes of Zhu et al. [24], of which 204 are among the 14,221 TSSs, were tested as an additional set.

Standard gene set enrichment analysis (GSEA) [38] assumes that every TSS is equally likely to rank high. However, in this case, how far a TSS can differ between the classes depends on its GC content, coverage and the depth of its dip in healthy plasma. Each set was therefore compared with 10,000 random sets of TSSs matched on these three properties. Two-sided empirical *p*-values were calculated as in Young et al. [39] and corrected with the Benjamini-Hochberg procedure. All TSSs were divided into 64 groups by the quartiles of the three properties in the 50 samples of the panel of normals. Each random set contains as many TSSs from each group as the gene set. A set whose TSSs have a higher Cohen’s *d* than those of its random sets is positively enriched, meaning that its promoters are more active (deeper coverage dips) in plasma of patients with cancer. For the genes of Zhu et al., the running enrichment score of a modified GSEA was compared with those of the same matched random sets.

#### 2.9.4. Classes of cell types

The C8 signatures were summarized into six classes by keywords in their names, without reference to the enrichment results: blood and immune cells, epithelial and parenchymal cells, neural and glial cells, stromal and mesenchymal cells, endothelial cells, and other (Supplementary Table S6). Signatures from bone marrow and cord blood were counted as blood and immune cells.

## 3. Results

### 3.1. Fragmentomic features at individual TSSs robustly capture chromatin structure in low-pass cfDNA

A data-driven locus selection framework requires a set of genomic regions with sufficient biological structure to support feature extraction at individual loci. We chose the transcription start site (TSS) of every protein-coding gene as a candidate feature. The nucleosome architecture at a TSS, a nucleosome-depleted region (NDR) flanked by positioned nucleosomes, reflects how active the gene is in the cells that release cfDNA [4, 21]. As the model treats each TSS as a feature on its own, we first tested whether this architecture is detectable at single TSSs at low coverage (mean ~2.48x for the cohort of Cristiano et al.). Aggregate coverage profiles of stably expressed housekeeping (HK) genes and unexpressed (PAU) genes recapitulated the abovementioned architecture, where expressed genes show a pronounced NDR while unexpressed genes display uniform coverage (Figure 1a). If the architecture is visible at single TSSs, the 10k bp window of an expressed gene should have a lower mean coverage than that of an unexpressed gene. Pooled over the 50 samples, the mean window coverage of single TSSs was indeed lower for HK than for PAU genes (Mann-Whitney U test, *p <* 0.001; Figure 1b). The nucleosome architecture signal is thus retained at individual TSSs. Given this structural consistency, we computed five fragmentomic features within each 10k bp TSS window (FSLR, FSLRC, GD, MWH, RCOV; Supplementary Table S3, Methods) [14, 3, 2, 24].

To assess whether this locus-level signal carries biological meaning, we correlated each feature with gene expression across the panel of normals, following Esfahani et al. [27]. We used all TSSs (n = 18,534, Methods) and the expression levels (log(TPM)) of a reference PBMC dataset (Figure 1c, Methods). Features that integrate signals across the full 10k bp window (FSLR/FSLRC and GD) exhibited strong correlations with expression. By contrast, features relying on narrower local signals within the window (RCOV and MWH) showed reduced sensitivity in this low-pass setting.

To further increase robustness, we applied the quality control filters (Section 2.2), retaining 14,226 TSSs. This increased the correlation of all features with gene expression (Figure 1c, hatched bars). A similar pattern was observed when the samples were downsampled to 1x, but not anymore at 0.1x (Supplementary Figure S6). At the sequencing depth of this cohort, the features thus reflect gene expression at single TSSs, supporting the use of each TSS as an individual feature.

### 3.2. Data-driven TSS analysis compared with predefined region sets and ichorCNA

#### 3.2.1. Benchmark design

We next assessed whether data-driven selection of informative loci improves cancer detection over predefined region sets. Our data-driven framework ranks all 14,221 filtered TSSs by their ability to separate healthy individuals from patients with cancer, and selects the top *k* loci. It then extracts a single fragmentomic feature per locus and trains a classifier on the resulting feature vectors (Section 2.3, Figure 2). The classification task was to distinguish healthy controls (*n* = 164) from patients with colorectal (CRC; *n* = 26) or breast cancer (BRCA; *n* = 54) in the Cristiano et al. cohort. These 80 patients form one class, called BRCA/CRC below. In the cross-validation results, the data-driven model refers to the outcome of this nested procedure: the model of each outer fold, evaluated once on its held-out fold, so that every participant has one out-of-fold prediction (Section 2.6).

**Fig. 2:**
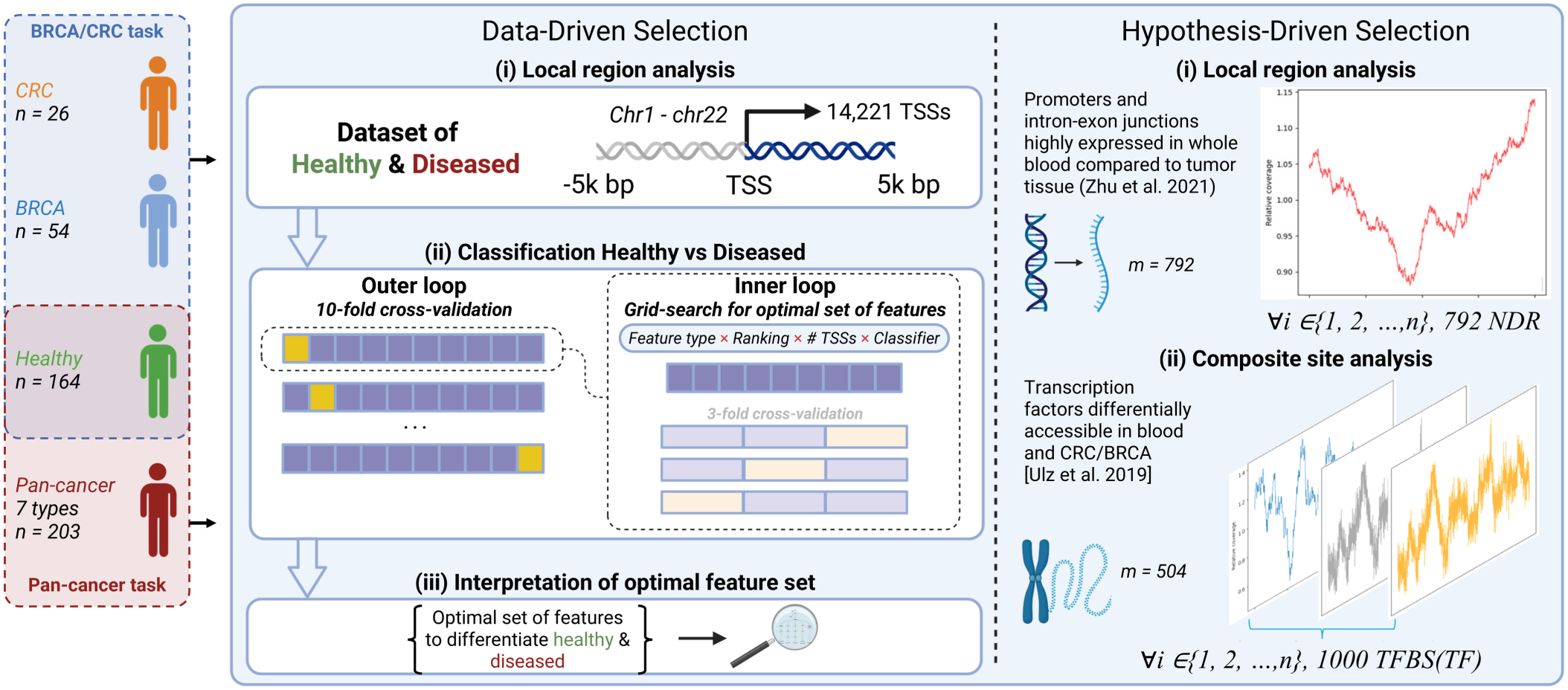
Method overview for binary classification using data-driven ROI. For the data-driven approach, nested cross-validation selects a set of TSSs of protein-coding genes, a fragmentomic feature and a classifier; the selected TSSs are then interpreted for their biological relevance. The hypothesis-driven baselines use the regions selected by Zhu et al. [24] and Ulz et al. [12]. Figure created with Biorender.com. *Alt text:* Diagram of the study design. Patients with breast or colorectal cancer form one classification task and patients with seven cancer types a second, and both tasks share the same healthy donors. A data-driven workflow selects transcription start sites, a feature and a classifier in nested cross-validation, next to two hypothesis-driven workflows built on blood-specific regions and on transcription factor binding sites.

CRC and BRCA are the two cancer types for which the hypothesis-driven region sets were defined a priori from external molecular datasets. The first, the RNA-seq–based region set of Zhu et al. [24], contains promoters and intron–exon junctions highly expressed in blood relative to solid tumor tissues. The second consists of the transcription factor binding sites (TFBSs) of the ATAC-seq–based approach by Ulz et al. [12], differentially accessible in CRC and BRCA relative to hematopoietic lineages. We also benchmarked against ichorCNA [7], a widely used copy number–based method that estimates tumor fraction. All methods used the same nested cross-validation (Section 2.6). For the two region sets, the regions and feature were fixed, and the classifier and its hyperparameters were tuned.

#### 3.2.2. Overall classification performance

For BRCA/CRC versus healthy, the data-driven model achieved a mean AUROC of 0.966 ± 0.029 over the ten outer folds of the nested cross-validation (Figure 3a). It exceeded the RNA-seq baseline (AUROC 0.666 ± 0.108) and ichorCNA (0.673 ± 0.079; both *p <* 0.001). It was also higher than the ATAC-seq approach (0.891 ± 0.096) in nine of ten folds. This difference was, however, not statistically significant (difference 0.075, 95% CI −0.038 to 0.188, *p* = 0.17; corrected resampled *t*-test, Holm-adjusted). Inall ten outer folds, the nested selection chose the GD feature and a linear support vector machine. It chose 10,000 TSSs in seven folds and all 14,221 in three, the two largest values of the search grid (Supplementary Table S7). At the 95% specificity threshold derived from the training folds, the data-driven model reached a sensitivity of 0.81 (65/80; 95% CI 0.71 to 0.89) at a specificity of 0.95 (156/164; 0.91 to 0.98; Figure 3b, Supplementary Table S7). At their own thresholds, the ATAC-seq baseline reached a sensitivity of 0.68 (54/80), the RNA-seq baseline 0.14 (11/80) and ichorCNA 0.30 (24/80), at specificities of 0.93, 0.90 and 0.96.

**Fig. 3:**
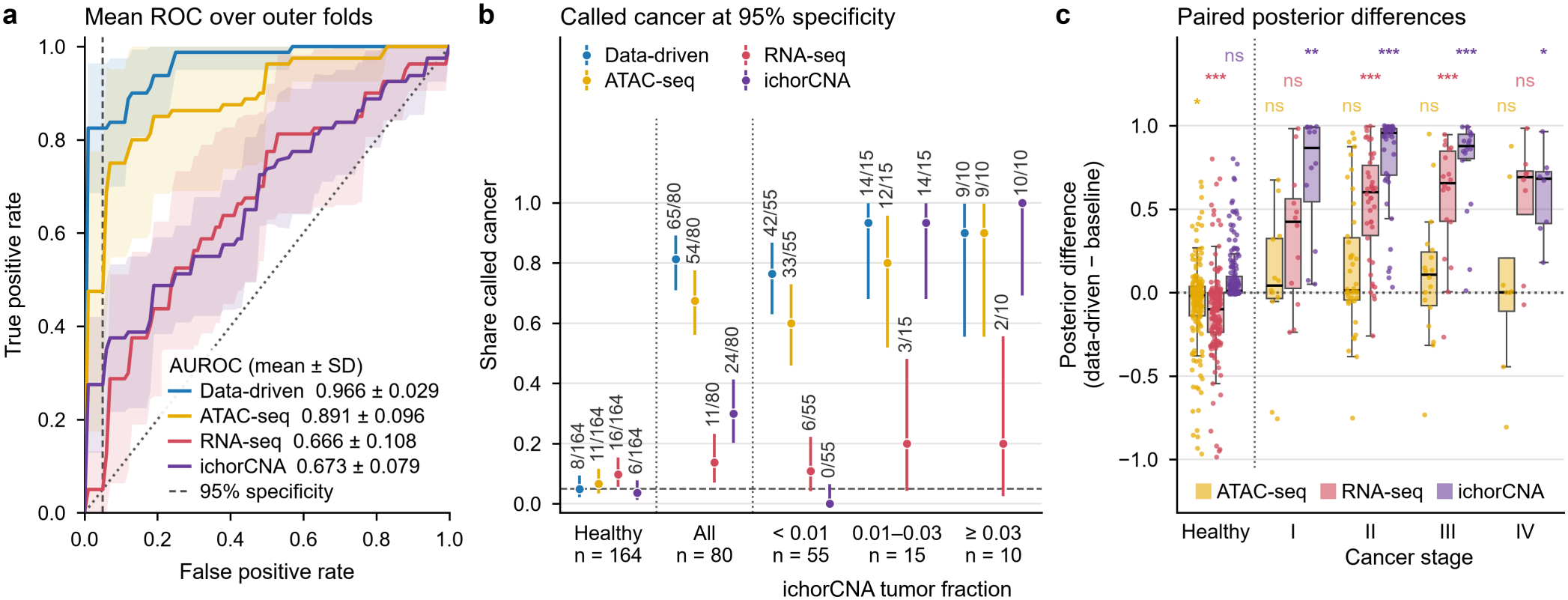
Classifier comparison. a) Nested CV performance on the binary classification task of healthy versus BRCA/CRC patients (breast and colorectal cancer together as one class) in the Cristiano et al. cohort. ROC curves are shown as the mean curve over the nested CV outer folds, with the shaded regions equaling one standard deviation. b) Share of participants called cancer by each method at its 95% specificity threshold, derived in each outer fold from the training samples only (for ichorCNA, from their tumor fractions): healthy donors, all cancers, and the cancers grouped by their ichorCNA tumor fraction. Out-of-fold calls pooled over the ten outer folds, with exact binomial (Clopper-Pearson) 95% confidence intervals; numbers give called / total, and the dashed line marks 5%. c) Paired difference per participant, by cancer stage: the posterior probability of the data-driven model minus that of the baseline (for ichorCNA, its tumor fraction). Points: single participants; boxes: median and interquartile range; whiskers: 1.5 times the interquartile range. Positive values denote a larger posterior probability for the data-driven model. Significance is indicated by a one-sided Wilcoxon signed-rank test to assert whether the data-driven model has significantly higher probabilities than the baseline model, Holm-adjusted over the 15 tests of the panel (ns, not significant; * *p <* 0.05; ** *p <* 0.01; *** *p <* 0.001). In the healthy class, we flipped the one-sided test to assess whether the data-driven model assigns lower posterior probabilities to healthy samples than each baseline. *Alt text:* Graphs labelled a to c comparing the data-driven model with three baseline methods for detecting breast and colorectal cancer. a, receiver operating characteristic curves, with the data-driven curve highest. b, dot plot of the share of participants called cancer at 95 percent specificity, between 4 and 10 percent of healthy donors and highest for the data-driven model among all cancers and at the lowest tumor fractions. c, box plots per cancer stage of the difference in predicted probability between the data-driven model and each baseline, mostly above zero for patients with cancer.

**Fig. 4:**
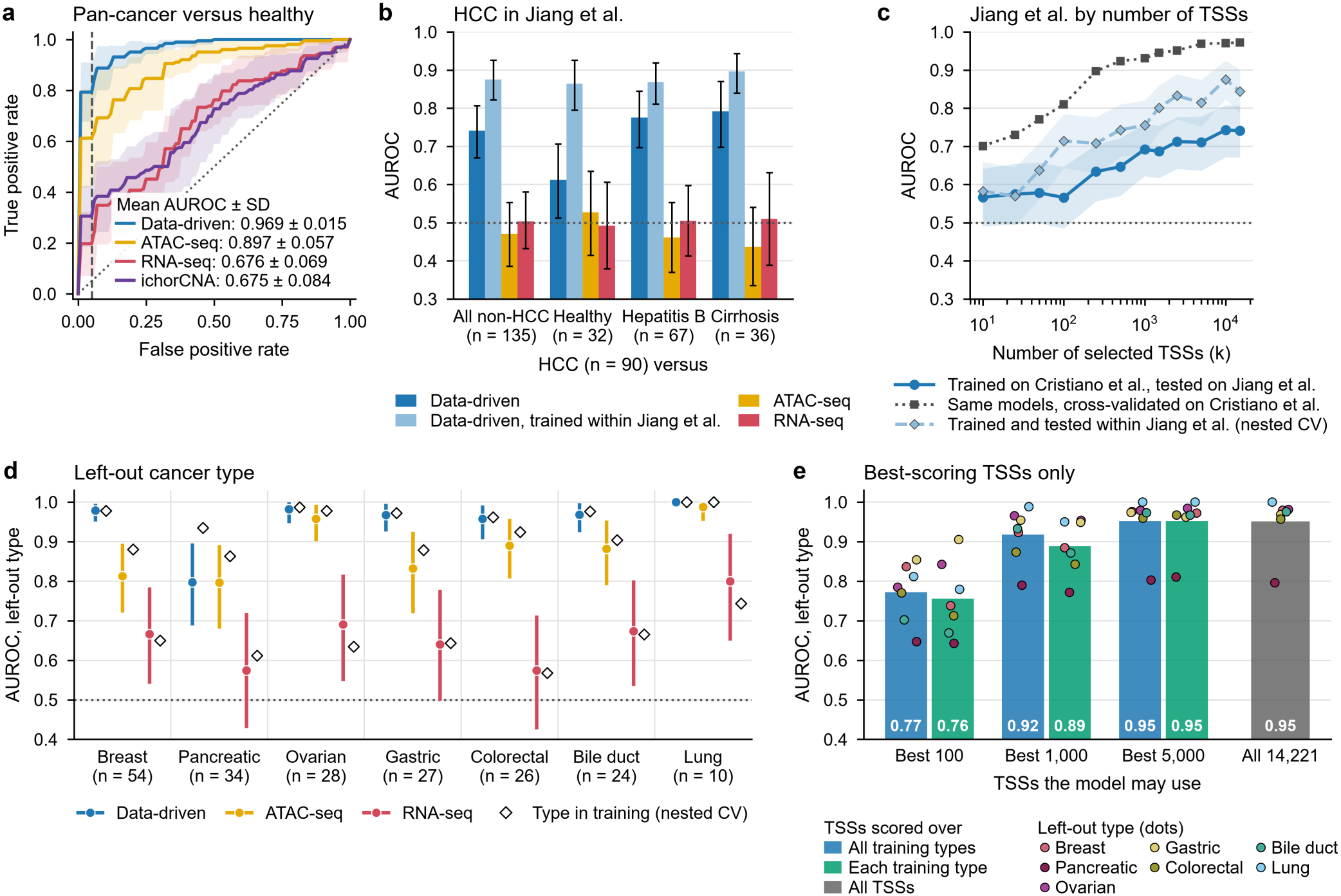
Performance in the PANCAN and generalization settings. a) Healthy versus all seven cancer types together, Cristiano et al. cohort. Mean ROC curve over the ten outer folds of the nested CV (shaded: *±* one standard deviation), with the mean AUROC *±* standard deviation in the legend. The dashed vertical line marks 95% specificity. b) AUROC on the Jiang et al. dataset for HCC (n =90) against all 135 non-HCC participants and against each comparison group, for the data-driven model trained on the Cristiano et al. cohort, the same procedure trained within the Jiang et al. cohort (nested CV, out-of-fold predictions), and the ATAC-seq and RNA-seq baselines trained on the Cristiano et al. cohort. Error bars: participant-level bootstrap 95% confidence intervals. c) AUROC against the number of selected TSSs (*k*): models trained on the Cristiano et al. cohort and tested on the Jiang et al. dataset (solid), the same models cross-validated on the Cristiano et al. cohort (dotted), and the same procedure trained and tested within the Jiang et al. dataset by nested cross-validation (dashed). Shaded: bootstrap 95% confidence intervals. d) Leave-one-cancer-type-out analysis. For each cancer type, the data-driven model and the ATAC-seq and RNA-seq baselines were selected and trained on 130 healthy donors and the six other cancer types, then applied once to the left-out type and 34 held-out healthy donors (number of patients in brackets). Circles: AUROC of each method (color as in the legend) on the left-out type against the held-out donors; lines: participant-level bootstrap 95% confidence intervals. Diamonds: the same patients and donors in the nested pan-cancer CV of the same method, where the type is in training. e) AUROC on the left-out cancer type when the configuration that all seven searches selected (GD, elastic-net logistic regression) is retrained with only the best-scoring 100, 1000 or 5,000 TSSs, or all of them. The TSSs are scored on the training samples by the absolute difference between group means, either over all training cancer types together (blue) or within each of the six training types with the ranks averaged (green); gray: all TSSs. Bars: mean of the seven left-out types; dots: single left-out types (key below the panel). Gray dotted lines at 0.5 (in a, the diagonal): chance. *Alt text:* Graphs labelled a to e on pan-cancer detection and generalization. a, receiver operating characteristic curves, with the data-driven model highest. b, bar chart for liver cancer in an independent dataset against each comparison group: the data-driven model trained on the first cohort is above chance, the same procedure trained within the independent dataset is higher, and both region-based baselines stay near chance. c, line chart in which performance on the independent dataset rises with more transcription start sites but stays below training within that dataset. d, dot plot per left-out cancer type, with the data-driven model near perfect except for pancreatic cancer and never below the baselines. e, bar chart in which fewer top-ranked start sites give lower performance.

#### 3.2.3. Performance across cancer stages and tumor fractions

We next examined whether our data-driven model assigns consistently higher posterior probabilities to patients with cancer than the baselines across disease stages. For every cancer sample, we computed the difference in posterior probabilities between the data-driven model and each baseline, stratified by cancer stage (Figure 3c). In all four stages, the data-driven model yielded significantly higher posterior probabilities than ichorCNA. Against the RNA-seq baseline, they were significantly higher in stages II and III. In stages I and IV, with 12 and 8 patients, the difference with the RNA-seq baseline was as large but not significant after adjustment. The advantage over the ATAC-seq baseline was smaller and not significant in any stage. For the healthy samples, the data-driven model assigned significantly lower predicted probabilities than the ATAC-seq and RNA-seq baselines. It thus separates the classes better through both the cancers and the healthy donors.

Detection was not limited to advanced disease or a high tumor fraction (Figure 3b; Supplementary Figure S7a-c). The AUROC of the data-driven model was 0.92 for stage I and 0.96 to 0.98 for stages II to IV. It detected 42 of the 55 cancers whose ichorCNA tumor fraction was below 0.01, within the range of healthy donors. The ATAC-seq baseline detected 33 of these, the RNA-seq baseline 6 and ichorCNA none (by design here).

### 3.3. Pan-cancer evaluation and generalization

#### 3.3.1. Pan-cancer classification performance

The hypothesis-driven baselines were originally designed and applied to BRCA and CRC detection, so it is uncertain how well they detect other cancer types. The data-driven framework, by contrast, selects its TSSs from the cfDNA of whatever cancers are in the training cohort. To test performance across cancer types, we repeated the full nested CV with all seven cancer types from the Cristiano et al. cohort (203 patients with cancer against 164 healthy controls; Section 2.6). All seven were included in both training and test folds. In this pan-cancer (PANCAN) task, the data-driven approach achieved a mean AUROC of 0.969 ± 0.015 (Figure 4a). It was higher than the RNA-seq baseline (0.676 ± 0.069) and ichorCNA (0.675 ± 0.083; both *p <* 0.001). It was also higher than the ATAC-seq baseline (0.897 ± 0.057) in nine of ten folds. Unlike in the BRCA/CRC task, this difference was statistically significant (difference 0.073, 95% CI 0.010 to 0.135, *p* = 0.028; corrected resampled *t*-test, Holm-adjusted over the three baselines). The baselines performed as well in the pan-cancer task as for BRCA/CRC, but none of them reached the data-driven model.

At the 95% specificity threshold derived from the training folds, the data-driven model reached a sensitivity of 0.81 (165/203; 95% CI 0.75 to 0.86) at a specificity of 0.98 (160/164; 0.94 to 0.99). As in the BRCA/CRC task, the nested selection chose GD and 10,000 or all 14,221 TSSs in every outer fold (five folds each). Different from that task, here it chose an elastic-net logistic regression classifier. Per cancer type, the AUROC ranged from 0.94 for pancreatic cancer to 1.00 for lung cancer. Performance by stage, cancer type and tumor fraction is given in Supplementary Figure S7d-g.

#### 3.3.2. Generalization to the Jiang et al. dataset

To evaluate generalization beyond the training cohort, we selected hyperparameters by cross-validation on the full Cristiano et al. cohort and retrained a single model on the entire cohort with that configuration. This model was applied without further tuning to the independent Jiang et al. dataset (Section 2.6), which consists of patients with hepatocellular carcinoma (HCC) and non-HCC participants. HCC is a cancer type that was not represented in the training set and the non-HCC class contains 32 healthy subjects, 67 patients with chronic hepatitis B and 36 with hepatitis B-associated cirrhosis [20]. The two liver disease groups differedsignificantly from the HCC patients and the healthy subjects in coverage and fragment GC content (two-sided Mann-Whitney U test, *p <* 0.001 for both; Supplementary Figure S2b), whereas the HCC patients and the healthy subjects did not differ in either (*p* = 0.57 and *p* = 0.09).

The data-driven model achieved an AUROC of 0.74 (95% CI 0.67-0.80) for HCC against all non-HCC participants. It reached0.78 against chronic hepatitis B, 0.79 against cirrhosis and 0.61 (95% CI 0.51-0.72) against the 32 healthy subjects (Figure 4b). The RNA-seq (0.50) and ATAC-seq (0.47) baselines did not outperform random guessing, both performing significantly worse than the data-driven model (difference 0.24, 95% CI 0.14 to 0.33, and 0.27, 0.18 to 0.37; DeLong test, Holm-adjusted *p <* 0.001 for both). Trained within the Jiang et al. cohort, the same nested procedure reached an AUROC of 0.88 (95% CI 0.82-0.93; Figure 4b). The signal is thus present in this cohort, but a model trained on the Cristiano et al. cohort captures only part of it.

At the threshold fixed on the Cristiano et al. cohort for 95% specificity, the data-driven model detected 39 of the 90 HCC patients (sensitivity 0.43; 95% CI 0.33 to 0.54). This came at a specificity of 0.89 (120/135; 0.82 to 0.94; Supplementary Table S7), a modest loss of specificity. The ATAC-seq baseline, by contrast,called nearly all participants cancer (specificity 0.10). Performance on the Jiang et al. dataset did not degrade as the number of selected TSSs increased (Figure 4c). The loss on the Jiang et al. dataset is thus not caused by the large number of TSSs that the nested selection chose.

#### 3.3.3. Generalization to unseen cancer types within the Cristiano et al. cohort

The lower performance on the Jiang et al. dataset could reflect differences between the cohorts, the unseen cancer type, or both. To separate the two, we performed a leave-one-cancer-type-out analysis within the Cristiano et al. cohort. Each method was selected and trained on six cancer types and 130 healthy donors. It was then applied once to the left-out type and 34 held-out healthy donors (Section 2.6). Six of the seven types reached an AUROC of 0.96 to 1.00 when left out (Figure 4d), the same range as in the pan-cancer task, where they were part of training. Pancreatic cancer was the exception, with 0.80 (95% CI 0.69 to 0.90) when left out against 0.94 in the pan-cancer task. Averaged over the seven left-out types, the data-driven model reached an AUROC of 0.95, against 0.88 for the ATAC-seq baseline (difference 0.07, 95% CI 0.02 to 0.12) and 0.66 for the RNA-seq baseline (0.29, 0.21to 0.37). It was better than the RNA-seq baseline for every type. Against the ATAC-seq baseline, it was higher for every type except pancreatic cancer (0.80 for both). The difference was significant for breast cancer (DeLong test, Holm-adjusted over the seven types, *p* = 0.002), interestingly the type whose ROIs partially inform the ATAC-seq model.

TSSs that score well in every training cancer type did not transfer better to an unseen type. When the model may use only the best-scoring 100, 1000 or 5,000 of them, it reached an AUROC of 0.76, 0.89 and 0.95, averaged over the seven left-out types. With all TSSs, it reached 0.95 (Figure 4e). All cancer types share the same technical setting in this analysis. The data-driven model thus captures a cancer-associated signal that is not restricted to the cancer types used during training. On average, its advantage over the hypothesis-driven baselines holds for cancer types it has not seen.

### 3.4. Performance is robust to covariates

Because an unseen cancer type cost the model little performance within one cohort, differences between the cohorts are the likely reason for the generalization gap on the Jiang et al. dataset. This motivated us to investigate whether the model is biased by technical and/or demographic properties of the samples rather than purely capturing the disease signal.

#### 3.4.1. Covariates within the Cristiano et al. cohort

We found that such covariates are influential in the Cristiano et al. cohort, as a classifier trained on coverage, fragment count, plasma volume, age and sex alone (selected by the same nested procedure) separated cancer from healthy participants at an AUROC of 0.88 (BRCA/CRC versus healthy) and 0.90 (pan-cancer versus healthy; Supplementary Figure S8a). We then asked whether, within the Cristiano et al. cohort, the model relies on these covariates. Adding the five covariates to the TSS features left its AUROC unchanged (0.966 to 0.966 and 0.969 to 0.972; Supplementary Figure S8b, c). The ranking of the TSSs by their difference in GD between cancer and healthy was the same after adjusting for the covariates (Spearman correlation of 0.96 and 0.99; Supplementary Figure S8d, e). Within healthy donors, the scores of the modelcorrelated at most weakly with any covariate (Spearman *ρ* ≤ 0.27; Supplementary Figure S8f). Within a class, the model ranked the samples differently from the classifier trained on the covariates, so the two do not use the same information (*ρ* between −0.12 and 0.07; Supplementary Figure S8g). From these results, we concludethat this cohort contains differences between cancer and healthy participants that are enough to reach very high classification performance, but that the data-driven model does not rely on this information.

#### 3.4.2. Covariates on the Jiang et al. dataset

To test whether such measures generalize to the Jiang et al. dataset, we trained a classifier in the same way on five measures that can be derived from the sequencing data of any cohort (coverage, fragment count, fragment length, GC content and sex). For pan-cancer versus healthy, this model achieved a cross-validated AUROC of 0.84 within the Cristiano et al. cohort (Supplementary Figure S8a), but 0.55 on the Jiang et al. dataset. There, the data-driven model reached 0.74 (Supplementary Figure S8h). Neither adding these measures to the model nor standardizing its features on the Jiang et al. dataset changed thatvalue. The data-driven model thus captures a discriminatory signal across cohorts that these measures do not.

### 3.5. Performance is robust to lower sequencing depth and smaller training cohorts

We then assessed the sensitivity of our data-driven approach to sequencing depth and cohort size. We downsampled every sample to 1x and to 0.1x (Supplementary Figure S9a-c, e-g). The models trained at full depth, applied unchanged to the downsampled samples, kept a mean AUROC of 0.936 at 1x for BRCA/CRC versus healthy, against 0.966 at full depth (not detectably lower; Holm-adjusted Wilcoxon *p* = 0.33), and of 0.938 for pan-cancer versus healthy, against 0.969 (lower in all ten outer folds; *p* = 0.020, the smallest value ten folds allow). Repeating the complete nested procedure at 1x did not improve on this (0.902 and 0.914; *p* = 0.33 and 0.21 against the full-depth models). At 0.1x, the AUROC fell to 0.734 and 0.686 for the full-depth models and to 0.649 and 0.662 after retraining, lower than every arm at full depth or 1x (*p* = 0.020 for each comparison). This is in line with the correlation of the features with gene expression, which was largely kept at 1x and much weaker at 0.1x (Supplementary Figure S6).

We then repeated the complete nested procedure on smaller subsets of the cohort. We compared the AUROC of a model trained on a subset with that of the model trained on the full cohort, on the same participants. With a paired participant-level bootstrap (Holm-adjusted over the subsets), it was not detectably lower down to 60% of the cohort for BRCA/CRC versus healthy and down to 80% for pan-cancer versus healthy. For pan-cancer versus healthy, the drop stayed small down to 60%, and it grew for smaller subsets (Supplementary Figure S9d, h). The approach thus keeps most of its performance at 1x coverage and with a moderately smaller training cohort, but markedly decreases in performance at 0.1x coverage or with a strongly reduced training cohort.

### 3.6. Discriminative signal is distributed across TSSs and benefits from modeling many TSSs individually

Nested cross-validation selected 10,000 or all 14,221 TSSs in every outer fold, for BRCA/CRC as well as for the pan-cancer task (Sections 3.2 and 3.3). To understand why so many TSSs were needed, we evaluated the four components of the framework. These are the number of selected TSSs (*k*), the ranking function, the classifier and the fragmentomic feature. We tested all combinations on the same ten outer folds, with only the hyperparameters of the classifier optimized in the inner loop (Section 2.6). This analysis was descriptive and did not influence model selection. Figure 5 shows the results for the pan-cancer task, with the AUROC of the fully nested procedure (0.969) as reference. The BRCA/CRC task showed similar patterns (Supplementary Figure S10a, b, d).

**Fig. 5:**
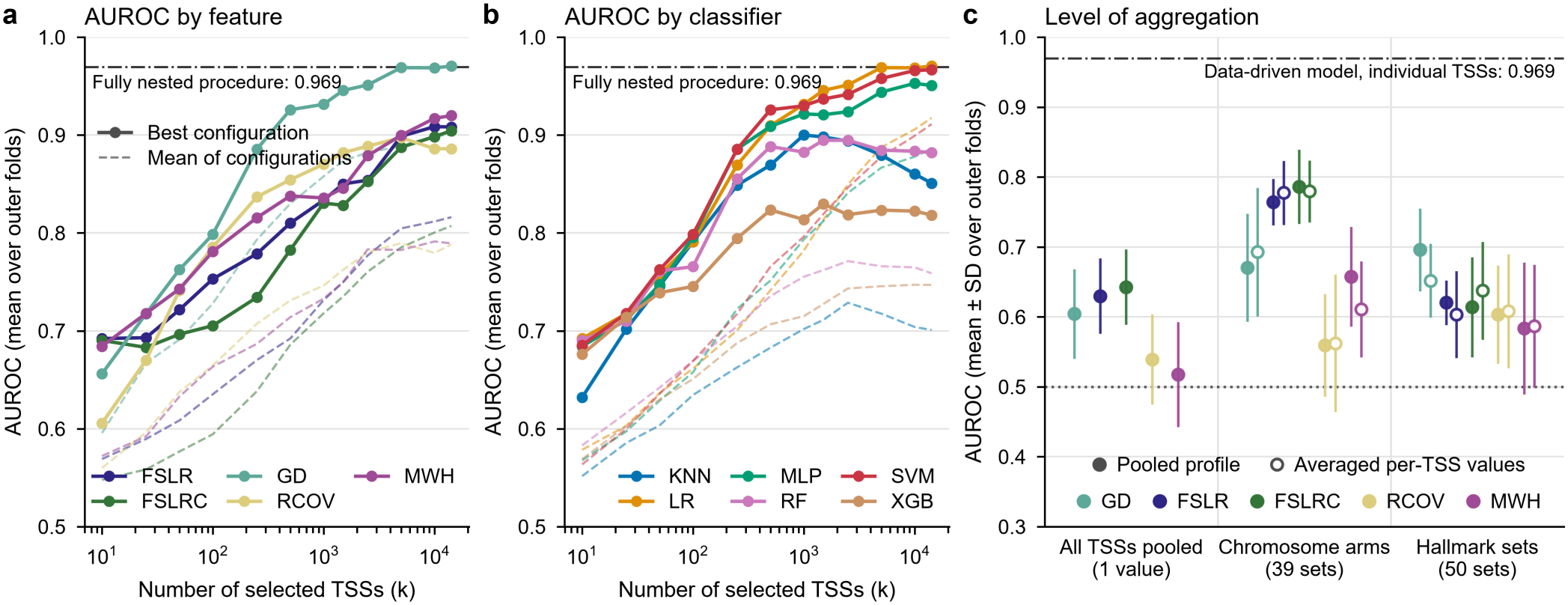
Analysis of the data-driven model for the pan-cancer task (healthy versus all seven cancer types). a) AUROC (mean over the ten outer folds) as a function of the number of selected TSSs (*k*), stratified by fragmentomic feature type. Solid lines: the best configuration at that *k*; dashed lines: the mean over all configurations. These curves are descriptive, because every configuration is scored on the outer test folds; the dash-dotted line marks the AUROC of the fully nested procedure, in which the configuration is selected within each outer fold (0.969). b) The same, stratified by classifier (KNN: K-Nearest Neighbors; LR: Logistic Regression; MLP: Multi-Layer Perceptron; RF: Random Forest; SVM: Support Vector Machine; XGB: XGBoost). c) AUROC by level of aggregation. Each fragmentomic feature was computed on all TSSs pooled (one value per sample), per chromosome arm (39 sets) and per Hallmark gene set (50 sets). Filled symbols: the feature computed once on the coverage profile (or fragment counts) pooled over the TSSs of a set. Open symbols: the feature computed at each TSS and then averaged over the TSSs of the set. Points: mean over the ten outer folds; error bars: one standard deviation; dash-dotted line: the data-driven model on individual TSSs (0.969); dotted line: chance. *Alt text:* Graphs labelled a to c analyzing the data-driven model for pan-cancer detection. a and b, line plots of performance against the number of transcription start sites, per feature and per classifier, rising with more sites for the best feature and for several classifiers. c, dot plot in which start sites aggregated into chromosome arms or gene sets perform well below the full model.

#### 3.6.1. Effect of the number of TSSs and chosen fragmentomic feature

Performance depended strongly on the number of selected TSSs (*k*). For the best-performing configurations, the mean AUROC increased as more TSSs were included and reached the fully nested value only from 5,000 TSSs on (Figure 5a, b). This gain was mainly driven by GD, the feature selected in every outer fold, which was above the four other features from 50 TSSs on and continued to improve as more TSSs were added, whereas none of the others exceeded 0.92 (Figure 5a). Together with its strong correlation with gene expression (Figure 1c), this suggests that the coveragedip at the TSS remains informative under low-pass conditions and benefits most from combining many TSSs.

As a control, we repeated the complete nested procedure with the class labels permuted once before the outer folds were drawn. The AUROC dropped to 0.51 for the pan-cancer task and 0.47 for BRCA/CRC (Supplementary Figure S10c). With the true labels, it was 0.969 and 0.966. This shows that the performance depends on the association between the TSS features and the true labels.

#### 3.6.2. Effect of classifier and TSS ranking

The benefit of more TSSs depended strongly on the classifier, and the AUROC gains at high *k* are carried by the linear models (Support Vector Machine and Logistic Regression), whereas the non-linear classifiers (Random Forest, K-Nearest Neighbors andXGBoost) plateau or decline beyond *k* ≈ 1000, with the Multi-Layer Perceptron in between (Figure 5b). This is consistent withknown limitations of these classifiers in high-dimensional, low-sample-size settings, and with the nested procedure selecting a linear classifier in every outer fold.

Ranking the TSSs also improved performance. All four ranking functions gave higher AUROCs than random selection of *k* TSSs for all *k <* 14,221, in both tasks. The Mann-Whitney U and mean-difference rankings performed best. With 1000 TSSs in the pan-cancer task, for example, they reached an AUROC of about 0.93, against 0.84 for randomly selected TSSs (Supplementary Figure S10e, f). Randomly selected TSSs thus carry useful signal.Ranking helps to find the most informative TSSs when only a subset is used.

#### 3.6.3. Hypothesis-driven baselines with a tuned number of regions

We also tested whether the hypothesis-driven baselines would gain from a tuned number of regions. With the number of regions selected in the inner loop, the ATAC-seq baseline rose slightly from 0.891 to 0.908 (BRCA/CRC versus healthy) and from 0.897 to 0.912 (pan-cancer versus healthy). The RNA-seq baseline changed little (Supplementary Figure S11). Both stayed below the data-driven model (0.966 and 0.969). Its higher AUROC is therefore not a consequence of tuning *k* for this model only.

#### 3.6.4. Locus-level features perform better than aggregated signals

Finally, we tested whether pooling coverage over many TSSs would serve these low-coverage samples better than features at individual TSSs. We computed each fragmentomic feature on the coverage pooled over all TSS windows of a sample, which gives a single value per sample. We did the same per chromosome arm (39 sets) and per Hallmark gene set (50 sets), pooled or as the average of the per-TSS values (Section 2.4). All aggregated models performed substantially worse than the model on individual TSSs (Figure 5c). Their AUROC was 0.52 to 0.64 for the single value, at most 0.79for the chromosome arms and at most 0.70 for the Hallmark gene sets. For individual TSSs, it was 0.969.

These results suggest that the discrimination does not arise from a uniform change across promoters. Instead, it depends on combining the signals of many TSSs that differ in their behavior. The tested aggregations, by chromosome arm or gene set, lose much of this information.

### 3.7. The distributed signal may reflect the cellular origin of cfDNA

We chose TSSs because cfDNA coverage around the TSS reflects the activity of its gene (Figure 1). The data-driven framework, owever, was optimized for cancer detection, not for identifying specific biological processes. We therefore examined which TSSs carry the signal, and whether it is shared between cancer types.

#### 3.7.1. TSS rankings are unstable at individual loci but shared across cancer types

We ranked the 14,221 TSSs by the standardized difference in GD, the depth of the coverage dip at the TSS, between patients with cancer and healthy donors (Cohen’s *d*). A positive difference here corresponds to a deeper coverage dip in cancer, i.e. a more active promoter. Between two disjoint halves of the cohort, the Spearman correlation of Cohen’s *d* was 0.28 (BRCA/CRC versus healthy) and 0.34 (pan-cancer versus healthy; Supplementary Figure S5a).

The 1,000 highest-ranked TSSs of the two halves overlapped hardly more than with permuted labels (Supplementary Figure S5b). The coefficients of the model were less stable still: even between outer folds, which share most of their training samples, they agreed less than Cohen’s *d* (Supplementary Figure S5c). No small set of marker TSSs can therefore be named, consistent with the model performing best with 10,000 or all TSSs (Section 3.6).

Despite this instability at single TSSs, the ranking as a whole was largely shared between cancer types. Relative to the agreement between two halves of the same type, the ranking of six of the seven types correlated with that of the others pooled at 0.57 to 0.92 (Figure 6a). Pancreatic cancer agreed least (0.38; 95% interval0.24 to 0.55). It was also the type detected worst when left out of training (Figure 4d). Pairs of single cancer types showed the same pattern (Supplementary Figure S5d, e). These results suggest that the discrimination rests on a broad pattern shared across cancer types, rather than on a small set of consistently ranked TSSs.

**Fig. 6:**
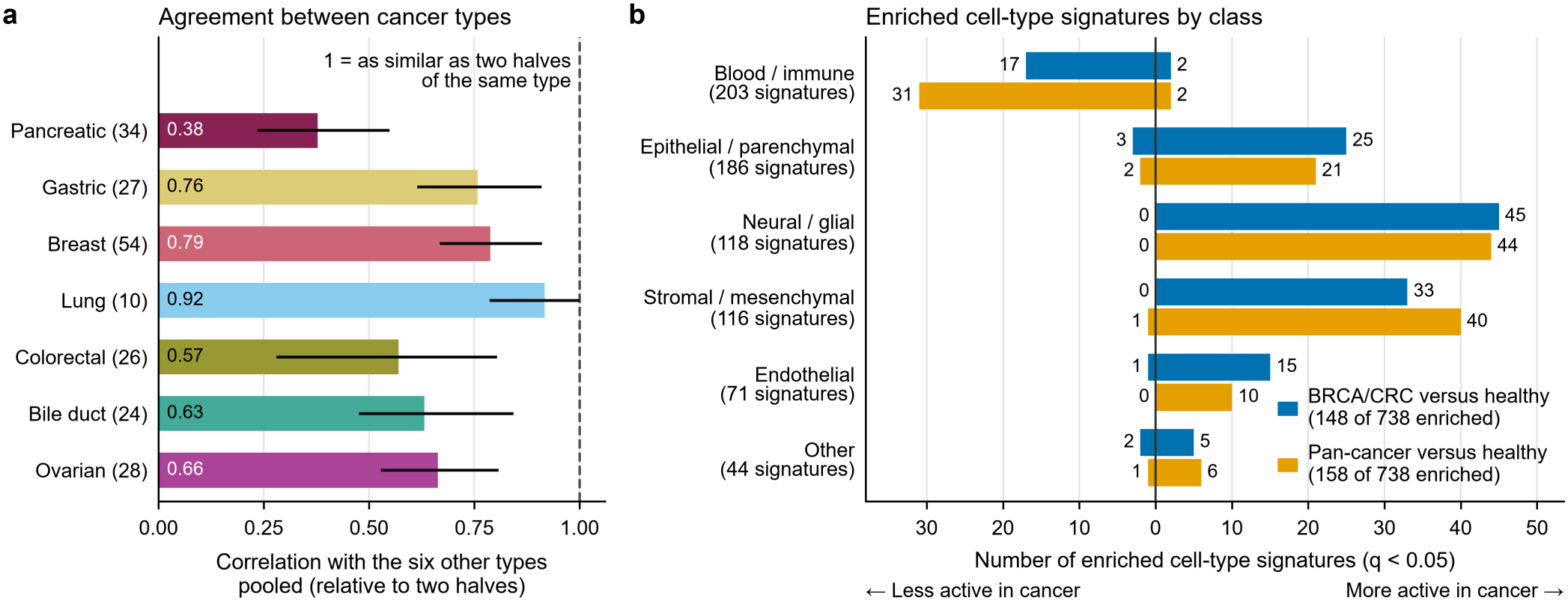
The ranking of the TSSs: agreement between cancer types and cell-type signatures. The TSSs are ranked by the standardized difference in dip depth (GD) between patients with cancer and healthy donors (Cohen’s *d*). a) Agreement of the ranking between cancer types: Spearman correlation of *d* between each cancer type and the six other types pooled, relative to the correlation between two halves of the same groups, so that 1 means as similar as two halves of the type itself (200 random splits; number of patients in brackets; colors as in Figure 4e). Lines: participant-level bootstrap 95% intervals (400 replicates). For lung cancer (10 patients), the two halves of the type did not agree at all in 101 of the replicates, so its interval has no upper bound; intervals are drawn between 0 and 1. b) Number of enriched cell-type signatures (MSigDB C8; 738 tested) per class of cell type, among the signatures that are negatively enriched (less active promoters; left) and positively enriched (more active promoters; right) in plasma of patients with cancer, for BRCA/CRC versus healthy (blue) and pan-cancer versus healthy (orange). Enrichment was tested along the ranking against 10,000 random sets of TSSs matched on dip depth, coverage and GC content in healthy plasma (*q <* 0.05). *Alt text:* Graphs labelled a and b on the ranking of transcription start sites in both tasks. a, bar chart of agreement between cancer types, lowest for pancreatic and highest for lung cancer. b, bar chart in which blood and immune cell signatures are mostly less active in cancer and the other cell types mostly more active.

#### 3.7.2. Cellular origins of differing TSSs

We tested gene sets along this ranking against random sets of TSSs matched on dip depth, coverage and GC content (Section 2.9). Of 738 tested cell-type signatures, 148 were enriched in BRCA/CRC versus healthy and 158 in pan-cancer versus healthy. Most that were negatively enriched, that is, whose promoters were less active in plasma of patients with cancer, were signatures of blood and immune cells (17 of 23 and 31 of 35). Nearly all that were positively enriched were signatures of other cell types (123 of 125 and 121 of 123; Figure 6b). The blood-specific promoter genes of Zhu et al.[24] were likewise negatively enriched (Supplementary Figure S12a, b). Few Hallmark sets and Reactome pathways, by contrast, were enriched (Supplementary Figure S12c, d). The signal thus likely reflects a change in the cellular origin of cfDNA in cancer, rather than a specific cancer pathway. In cancer, there is relatively less cfDNA from blood cells and more from other cells.

## 4. Discussion

In this study, we show that fragmentomic features measured at individual transcription start sites in low-pass whole-genome sequencing of cfDNA can distinguish patients with cancer from healthy individuals. Rather than relying on predefined, disease-specific genomic annotations, our data-driven framework starts from the TSSs of all protein-coding genes and ranks and integrates their fragmentation signal directly from the cfDNA data, within a nested cross-validation scheme. In both the BRCA/CRC and the pan-cancer task, it reached a higher AUROC than the RNA-seq baseline and ichorCNA; against the ATAC-seq baseline, the difference was statistically significant in the pan-cancer task but not in the BRCA/CRC task. Every outer fold selected 10,000 or all filtered TSSs, which indicates that the discriminative signal is distributed across loci rather than confined to an identifiable subset. The model also detected cancer types that were left out of training, and part of its signal carried over to the Jiang et al. dataset, which was generated with a different protocol and on which both hypothesis-driven baselines performed at chance.

One of our main findings is that the signal for cancer detectionis spread over many TSSs rather than concentrated in a few strongly informative loci. Performance kept increasing as more TSSs were included, and although the ranking of single TSSs was poorly reproducible, the ranking as a whole was largely shared between cancer types. Together, this suggests that cancer may be associated with a widespread change in cfDNA fragmentation, much of which is common to the cancer types we studied.

The cell-type signatures offer a possible biological explanation for this pattern. In plasma of patients with cancer, signatures of blood and immune cells were negatively enriched and those of other cell types positively enriched. This is consistent with a shift in the cellular origin of cfDNA, with relatively less DNA coming from blood cells and more from other cell types. Such a shift would change many promoters a little rather than a few strongly, which could explain both why the signal is spread over thousands of TSSs and why it is similar between cancer types. It may reflect a systemic change that accompanies cancer. In addition to the DNA released by the tumor itself, cancer may alter how much cfDNA different cells release, for example through tissue damage and the death of non-hematopoietic cells. Cell-type signatures in cfDNA fragmentation profiles have been linked to disease before [8]. A change of this kind could also explain why the model detected cancers with a low estimated tumor fraction, as well as cancer types left out of training. Alternatively, some of these tumors may release DNA without large copy-number alterations, which ichorCNA cannot detect.

This interpretation is consistent with our findings, but we cannot provide direct evidence for it since the depth of the coverage dip at a TSS does not directly measure how much cfDNA each cell type releases. Similar alterations have also been observed in inflammatory conditions independent of malignancy [16]. Also, while measured covariates did not explain the model, acquisition site and processing batch were not recorded for this cohort, so patients and healthy donors may still differ with respect to these variables. That few Hallmark sets and Reactome pathways were enriched argues against a signal dominated by a single cancer pathway, although it does not exclude contributions of specific processes. Studies that include non-malignant inflammatory conditions would show how much of the signal is specific to cancer. Our results also help to explain the differences between the data-driven framework and the hypothesis-driven baselines. Both baselines restrict the analysis to regions selected in advance from external molecular data. This can increase sensitivity at informative sites, but may miss changes in cfDNA fragmentation outside the selected regions.

Both hypothesis-driven baselines were originally designed for BRCA and CRC detection. The ATAC-seq approach nonetheless kept its performance in the pan-cancer setting, and was well above the RNA-seq baseline in both tasks. Its pan-cancer performance may suggest that the selected TFs play regulatory roles beyond the cancer types for which they were originally identified. Alternatively, aggregating the coverage over the 1000 binding sites of each TF may reduce sampling noise and coverage artifacts at low depth. Its performance falling to chance on the Jiang et al. dataset, however, may suggest that these coverage signals are cohort-specific.

Our aggregation experiments point to a further difference between the approaches. The TSSs lost much of their discriminatory power when pooled into one value per sample, per chromosome arm or per Hallmark gene set (Figure 5c), whereas the ATAC-seq approach preserves a per-TF resolution (497 features). It thereby retains part of the heterogeneity across loci that our analyses suggest is important for classification. Averaging may also suit its regions better since the binding sites of one TF were selected to behave alike, whereas the TSSs of a chromosome arm or a gene set are heterogeneous, so that pooling them removes the differences a classifier can use. The baseline also performed better in our BRCA/CRC task (AUROC 0.89) than in the original report of Ulz et al. [12], who give an accuracy of approximately 0.70, possibly because their evaluation focused on early-stage cancers with lower circulating tumor DNA fractions.

The RNA-seq baseline performed substantially worse than theATAC-seq approach in both tasks, despite targeting blood-specific regions rather than any single cancer type. That premise holds in our data since the TSSs of the blood-specific promoter genes of Zhu et al. [24] were less active in plasma of patients with cancer in both tasks (Supplementary Figure S12a, b). This is further corroborated by the negative enrichment of signatures of blood and immune cells, whereas signatures of other cell types were positively enriched. The lower performance of the baseline therefore points less to a wrong premise than to a signal that extends beyond the hematopoietic lineage. Capturing changes in the amount of cfDNA derived from blood cells alone may be insufficient. The regulatory dynamics at TFBSs may carry additional discriminatory information, as may the lower noise of the per-TF aggregation and the different features of the two baselines (RCOV and TFA).

With 14,221 TSSs and five features derived from them, the search space is large compared with the number of samples, the setting in which overfitting is a concern. We therefore made every model choice (feature, number of TSSs, ranking function and classifier) within a fully nested cross-validation, separate from performance evaluation, so that the reported performance is not inflated by model selection. Estimates from single test folds, in particular the sensitivity at 95% specificity, are more variable because the folds are small, which is why we report them pooled over the folds with confidence intervals.

Performance collapsed to random guessing when the class labels were shuffled, so the model captures a signal that is tied to the true labels. More importantly, the larger feature space did not hinder generalization, neither within the Cristiano et al. cohort nor on the Jiang et al. dataset: a model restricted to its best-scoring TSSs did not transfer better to a left-out cancer type, and performance on the Jiang et al. dataset did not degrade as the number of TSSs increased (Figure 4c, e). A positive relationship between feature dimensionality and classification performance has also been observed independently in a related cfDNA fragmentation analysis [40], which suggests that it is a broader property of distributed fragmentomic signals rather than an artifact of our particular experimental setup.

The benefit of many TSSs was most pronounced in thelinear models since they kept improving as more TSSs were included, whereas most non-linear classifiers plateaued or declined. Linear models may, therefore, suit the integration of distributed fragmentomic signals when the features far outnumber the samples. Together, these observations support combining many individually weak signals for cancer detection. Larger training cohorts may allow non-linear classifiers to learn from many TSSs with less risk of overfitting, and to model interactions between them that a linear model cannot.

Sequencing depth and cohort size also affected performance. Highly localized features (MWH and RCOV, which use a few hundred base pairs around the TSS) are more sensitive to sampling variability under low-pass sequencing than features that integrate information across the whole 10 kb window (FSLR, FSLRC and GD). Most of the discrimination ability was retained at 1x coverage, but it deteriorated substantially at 0.1x, where the correlation of the features with gene expression was also much weaker. When the size of the training set was reduced, performance was similar on the larger subsets but lower on the smaller ones, so a model that learns both its TSSs and its classifier from the data does need a sufficiently large training cohort. Still, trained on 40% of the total pan-cancer cohort, our model reached a numerically higher AUROC than the ATAC-seq baseline on the full cohort. Additionally, even larger training cohorts may also help obtain more stable estimates of the underlying biological signal, such as TSS rankings.

In the Cristiano et al. cohort, technical and demographiccovariates alone separated cancer from healthy participants at an AUROC roughly similar to the best hypothesis-driven baseline. This reemphasizes the known limitations of drawing conclusions by only applying cross-validation within one cohort [41, 42]. It is thus important to report such baselines next to fragmentomic models. Our additional experiments showed that adding these covariates to the TSS features did not change the data-driven model’s performance. Furthermore, the covariates were at most weakly related to its scores, and adjusting for them did not change the ranking of the TSSs. All these point to the conclusion that the model does not rely on these technical and demographic factors for its predictions. More importantly, a classifier trained on these features did not generalize to the Jiang et al. dataset (from an AUROC of 0.84 to 0.55), whereas the data-driven model reached 0.74. Some covariates, such as age, are themselves associated with cancer risk, so a covariate classifier that separates the groups well does not by itself indicate confounding. We nevertheless recommend comparing fragmentomic models with such a simple baseline classifier and testing both on an independent dataset.

On the Jiang et al. dataset, the data-driven model and both hypothesis-driven baselines showed a notable generalization gap, although the data-driven model kept part of its signal where both hypothesis-driven baselines failed to outperform random guessing. This gap likely reflects multiple factors. The Jiang et al. dataset differs from the Cristiano et al. cohort in read length, cfDNA extraction and library preparation (Supplementary Table S2), which introduces technical variability that affects fragmentation-based methods [43]. Because we kept the Jiang et al. dataset entirely out of training as an independent test, the model had not seen such differences between studies and could not learn to correct for them.

In addition, HCC is not among the cancer types in the training cohort. It arises from hepatocytes, which are, after endothelial cells, the largest non-hematopoietic source of cfDNA in healthy plasma [44]. If the model partly detects a shift from blood cells toward other cells, DNA released by an HCC may stand out less against this background, an effect that we cannot separate from the technical differences between the cohorts. The performance drop was observed for all three models, whereas an unseen cancer type showed little loss in performance within the Cristiano et al. cohort for six of the seven types. This suggests that cohort-level differences, rather than the number of TSSs or the absence of the cancer type from training, contribute most to the gap. The approach itself, however, is not tied to one cohort, since the same procedure trained within the Jiang et al. dataset reached an AUROC of 0.88. The higher AUROCs against the two liver disease groups than against the healthy subjects should be read with caution, as these groups differed from the HCC patients in coverage and fragment GC content, which may ease their separation from HCC independently of disease.

Our findings are also specific to TSS-based features. TSSsprovide a stereotypical chromatin structure with an NDR and flanking nucleosomes, which supports features that quantify depletion and oscillatory patterns [4, 21], whereas other regulatory elements (e.g., enhancers or TFBSs) may exhibit different fragmentation and coverage patterns and may therefore benefit from alternative feature definitions [13, 12, 2]. Whether genome-wide bins or enhancer regions would carry more signal was not tested. Our conclusions concern how regions are chosen, not whether TSSs are the best region set.

In summary, a data-driven framework that selects TSSs directly from cfDNA sequencing data reached a higher AUROC than hypothesis-driven region sets and ichorCNA, also for cancer types left out of training. Its signal is spread over thousands of TSSs, is largely shared between cancer types and is consistent with changes in the cellular origin of cfDNA. On an independently generated dataset it kept part of its signal where the hypothesis-driven baselines did not, and validation in larger cohorts processed independently of the training data is the next step.

## Data availability

The fragment-level data of both cohorts, Cristiano et al. [3] and Jiang et al. [20], are publicly available from FinaleDB (http://finaledb.research.cchmc.org/) [17]. Every FinaleDB record of the Cristiano et al. cohort, with its role in this study and thereason for any exclusion, is listed in Supplementary Table S1. The code to recreate the features from these data, train and evaluate all models, and draw every figure is available at https://github.com/DelftBioinformaticsLab/comp and archived at Zenodo [45] (release v1.0.0, https://doi.org/10.5281/zenodo.24462137; all versions: doi:10.5281/zenodo.24462136). The source data of every figure is included in the repository.

## Supplementary data

Supplementary Data are available at NAR Genomics and Bioinformatics Online.

## Author contributions statement

All authors meet the criteria for authorship.

- Conceptualization: BP, SM, SW, MR
- Methodology: BP, SM, SW, MR
- Software: BP
- Formal analysis: BP, SM
- Investigation: BP, SM
- Resources: SM, SW, MR
- Data Curation: BP
- Writing - Original Draft: BP
- Writing - Review & Editing: BP, SM, SW, MR
- Visualization: BP
- Supervision: SM, SW, MR
- Funding acquisition: SW, SM

## Acknowledgments

We express our gratitude to the past and present members of the Delft Bioinformatics Lab for their feedback on intermediate states of the work, conceptual discussions and code reviews. The authors used Claude (Anthropic; Opus 5, 5.5) to help write and refactor analysis and figure code, to check reported numbers against the result files, and to suggest edits to the text. The authors directed this use, reviewed and verified all output, and take full responsibility for the content of this article.

## Supplementary Material

**Table S1.** Participants of the Cristiano et al. cohort (separate Excel file, Supplementary_Table_S1.xlsx). One row per FinaleDB sequencing record (534 records of 473 individuals): role in this study, reason for any exclusion, task membership and, per task, the outer cross-validation fold, the out-of-fold probability of the data-driven model with its 95%-specificity threshold and call, the out-of-fold scores of the ATAC-seq and RNA-seq baselines, and the ichorCNA tumor fraction. A README sheet defines every column.

|  | Cristiano et al. 2019 (DELFI) | Jiang et al. 2015 |
|---|---|---|
| Data source | FinaleDB, hg38 fragment files | FinaleDB, hg38 fragment files |
| Samples analysed | Healthy 164, BRCA/CRC 80, other cancer types 123 | Healthy 32, chronic hepatitis B 67, HBV cirrhosis 36, HCC 90 |
| Comparison group | Healthy individuals | 135 non-HCC: 32 healthy, 67 chronic hepatitis B carriers, 36 HBV-associated cirrhosis |
| Blood collection and plasma processing | EDTA tubes, processed within 1 day; 800 g 10 min at 4 °C, then 18,000 g at room temperature; plasma at −80 °C | EDTA tubes; 1,600 g 10 min at 4 °C, then 16,000 g 10 min at 4 °C; plasma at −80 °C |
| cfDNA extraction | Qiagen Circulating Nucleic Acids Kit | QIAamp DSP DNA Blood Mini Kit (3–4.8 mL plasma) |
| Library preparation | NEBNext DNA Library Prep Kit for Illumina (modified), Phusion Hot Start amplification; PCR cycles not reported | KAPA Library Preparation Kit, 14 PCR cycles (KAPA HiFi HotStart ReadyMix) |
| Size selection | None reported (AMPure XP clean-up only) | None reported |
| Sequencing | HiSeq 2000, 100 bp paired-end | HiSeq 2000, 75 bp paired-end |
| Age (years) | Healthy 54 [50–75], BRCA/CRC 59 [36–83], other cancer types 64 [14–83] (not reported for 6 bile duct cancer, 5 colorectal cancer, 4 breast cancer) | Not reported |
| Female, inferred from the reads (%) | Healthy 68, BRCA/CRC 84, other cancer types 57 | Healthy 47, chronic hepatitis B 33, HBV cirrhosis 31, HCC 17 |
| Genome-wide coverage (×) | 2.42 [1.06–5.25] | 2.02 [1.20–4.84] |
| Fragments per sample (millions) | 39.9 [17.5–85.6] | 33.3 [19.6–79.0] |
| Mean fragment length (bp) | 165.0 [153.5–171.5] | 169.9 [153.8–177.9] |
| Modal fragment length (bp) | 166 [164–168] | 166 [165–175] |
| Short/long fragment ratio (100–150 / 151–220 bp) | 0.250 [0.105–0.674] | 0.140 [0.063–0.806] |
| Mean fragment GC (%) | 38.3 [35.8–45.7] | 41.7 [38.2–44.4] |
| ichorCNA tumor fraction, cancer samples | 0.008 [0.000–0.577] | 0.024 [0.011–0.368] |

**Table S2.** Characteristics of the Cristiano et al. (development) and Jiang et al. (additional) cohorts. Rows on sample collection, cfDNA extraction, library preparation and sequencing are taken from the Methods of both studies. Where applicable, values are the median [minimum-maximum] over the samples analysed (367 Cristiano and 225 Jiang samples), computed with our pipeline: coverage from the aligned reads, fragment-length statistics within the 100–220 bp window used for feature extraction, sex inferred from the reads on chromosomes X and Y, and ichorCNA tumor fraction in the cancer samples only. Age is the age at diagnosis and is not reported for the Jiang et al. cohort. They report a median tumor fraction of 2.1% in HCC with their own method. BRCA, breast cancer; CRC, colorectal cancer; HBV, hepatitis B virus; HCC, hepatocellular carcinoma.

| **Feature** | **Description** |
|---|---|
| Fragment Short/Long Ratio (FSLR) | Calculated as: $\log_2(\frac{\text{short}}{\text{long}})$, where *short* is the number of fragments between 100–150 bp and *long* is the number of fragments between 151–220 bp. |
| Fragment Short/Long Ratio (FSLRC) | Calculated as: $\log_2(\frac{\texttt{Coverage of Short Fragments}}{\texttt{Coverage of Long Fragments}})$. Using coverage instead of absolute counts is how the FSLR was originally introduced by Cristiano et al. |
| Griffin Difference (GD) | The coverage is normalized to a mean coverage of 1 across the 10 kb window. Then, the Griffin difference is calculated as the mean coverage of the 10 kb window outside the −990 to 990 bp window around the TSS minus the mean coverage inside that window, so that a deeper coverage dip at the TSS gives a larger GD. |
| Max Wave Height (MWH) | The coverage is normalized to a mean coverage of 1 across the 10k bp window. The MWH is calculated in the window from -120 bp to +195 bp relative to the TSS, as the absolute difference between the minimum coverage in the promoter region (-120 bp to +30 bp) and the maximum coverage at the first nucleosome in the gene body (+31 bp to +195 bp). |
| Relative NDR Coverage (RCOV) | Calculated by dividing the mean raw coverage around the ROI (-150 bp to 50 bp relative to the TSS) by the mean coverage of the 2000 to 1000 bp flanks up- and downstream of the ROI. These flanks serve as a normalization factor. |
| Transcription Factor Accessibility (TFA) | The coverage across a 2k bp window centered around all the top 1000 TFBS of one TF is first aggregated. This aggregated coverage is normalized to have a mean coverage of 1. Two Savitzky-Golay filters were applied for detrending. One filter captures the low-frequency signal of the coverage profile (third-order polynomial and window size of 1001) and one for the high-frequency signal (third-order polynomial and window size of 51). The high-frequency signal was adjusted by dividing by the low-frequency signal. The derived feature is the amplitude of this adjusted signal, as proxy of the TF accessibility, defined as the maximum value minus the minimum value. |

**Table S3.**
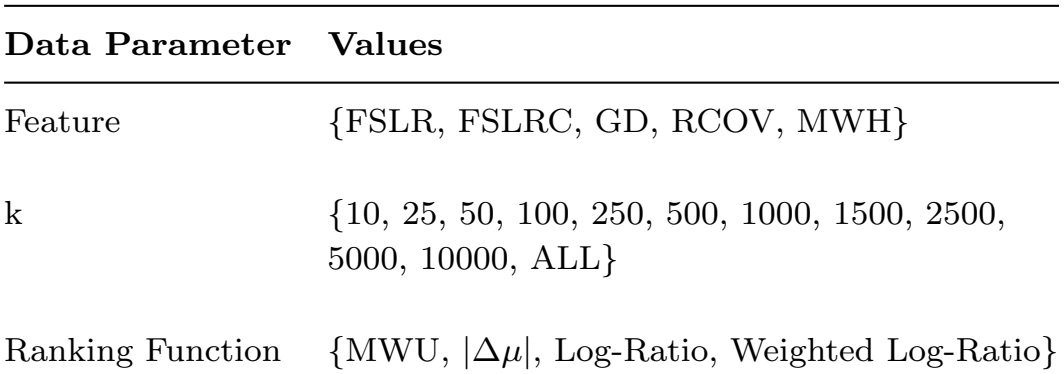
Description of features of the cfDNA fragments calculated at specific windows of interest.

| **Data Parameter** | **Values** |
|---|---|
| Feature | {FSLR, FSLRC, GD, RCOV, MWH} |
| k | {10, 25, 50, 100, 250, 500, 1000, 1500, 2500, 5000, 10000, ALL} |
| Ranking Function | {MWU, \|Δμ\|, Log-Ratio, Weighted Log-Ratio} |

**Table S4.** Data Parameters.

| Classifier | Hyperparameters |
|---|---|
| Support Vector Machine | *C*: 10<sup>−3</sup> to 10<sup>2</sup> (6 values, log-spaced)<br>Kernel: {linear, rbf}<br>Gamma: {scale, auto} |
| Logistic Regression | *C*: 10<sup>−3</sup> to 10<sup>1</sup> (5 values, log-spaced)<br>Penalty: elasticnet<br>Solver: saga<br>*l1_ratio*: {0.1, 0.5, 0.9} |
| Random Forest | *n_estimators*: {200, 500}<br>*max_depth*: {5, 10, None}<br>*min_samples_leaf*: {5, 10}<br>*max_features*: {sqrt, log2} |
| XGBoost | *n_estimators*: {100, 300}<br>*learning_rate*: {0.05, 0.1}<br>*max_depth*: {2, 3, 4}<br>*colsample_bytree*: {0.3, 1.0}<br>*subsample*: 0.8 |
| K-Nearest Neighbors | *n_neighbors*: {5, 7, 11, 15, 21}<br>*weights*: {uniform, distance}<br>Metric: {euclidean, manhattan} |
| Multi-Layer Perceptron | Hidden Layers: {(100), (200), (100, 50), (256, 128), (512, 256), (256, 128, 64)}<br>*α*: 10<sup>−4</sup> to 10<sup>−1</sup> (4 values, log-spaced)<br>Learning Rate: {0.001, 0.01}<br>*max_iter*: 2000 |

**Table S5.** Classifiers and their Hyperparameter Grids

Outer and inner folds were stratified with a fixed random seed (42), which was also used for all classifiers. No class weighting was applied. Scores are the classifiers’ native probabilities, with Platt scaling for the Support Vector Machine only. Features were standardized on the training samples of each fold; undefined feature values were set to zero after the per-sample z-score.

Hyperparameters, feature set and *k* were selected by the mean AUROC over the inner folds; among configurations with an equal inner-loop AUROC, the first hyperparameter setting of the grid and the smallest *k* were kept. The Logistic Regression selected in every outer fold of the pan-cancer task converged within its iteration limit (at most 4,126 of 10,000 iterations). The configuration selected inevery outer fold is listed in Supplementary Table S7.

**Table S6. Classes and enrichment of the MSigDB C8 cell-type signatures** (separate Excel file, Supplementary Table S6.xlsx). For each of the 738 signatures: the class of cell type assigned from its name, the number of TSSs, and per task the enrichment *z*, the *q* value and the direction against look-alike TSSs.

**Table S7. Selected configuration and confusion counts per outer fold** (separate Excel file, Supplementary Table S7.xlsx). Part A: per task, method and outer fold, the configuration selected in the inner cross-validation, the inner and outer AUROC, the 95%-specificity threshold set on the training partition, and the counts on the test fold. Part B: the counts pooled over the ten test folds with exact binomial 95% confidence intervals, and the Jiang et al. cohort at the threshold fixed on the Cristiano et al. cohort, for HCC against all non-HCC participants and against each comparison group.

**Fig. S1:**
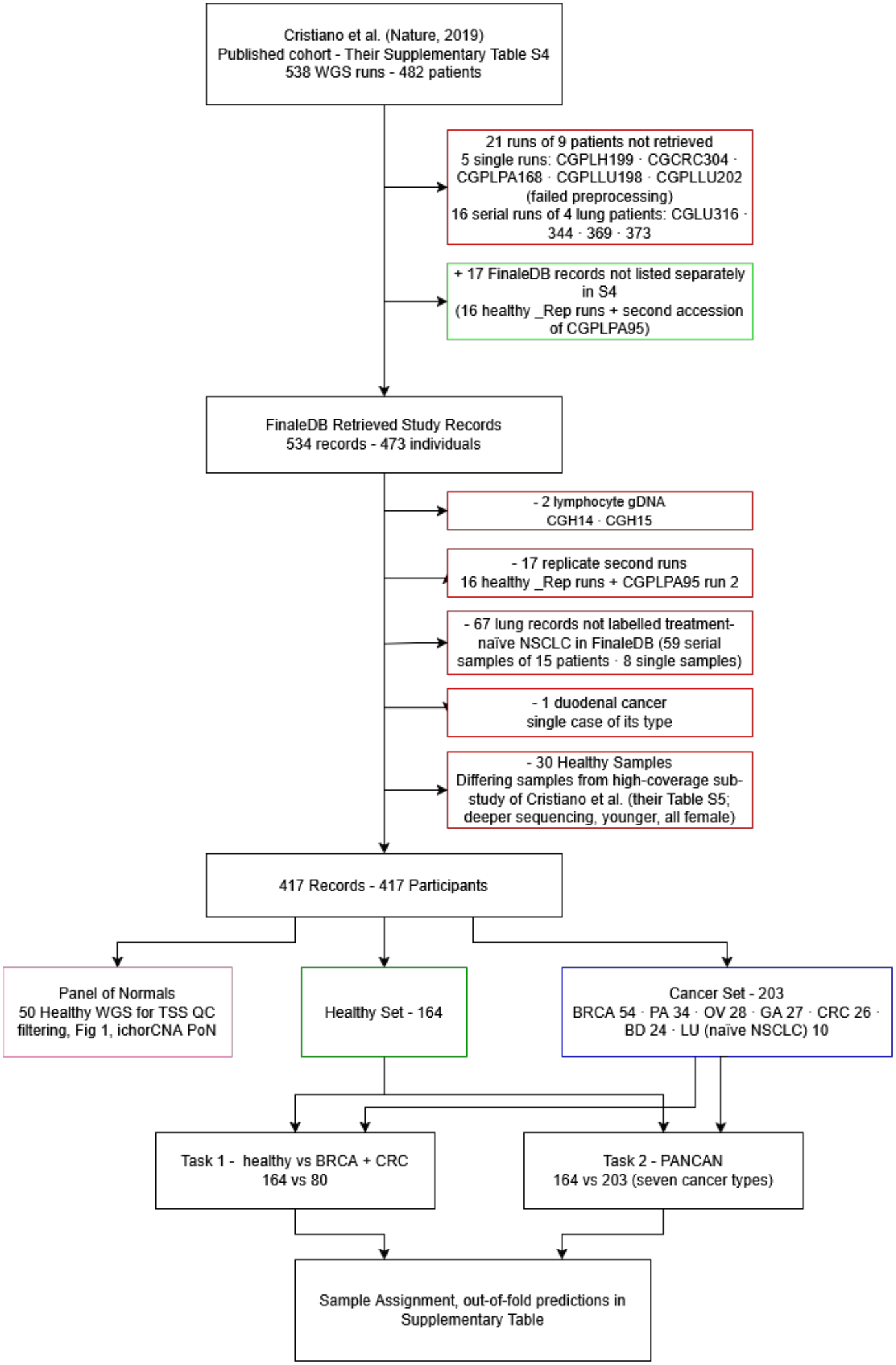
Sample flow of the Cristiano et al. cohort, from the 538 sequencing runs in their Supplementary Table S4 to the two classification tasks. Five runs failed preprocessing; the 16 serial runs of four lung patients with cancer that are not in FinaleDB would have been excluded as serial samples. FinaleDB holds 17 records that Table S4 does not list separately (green); these are second sequencing runs and are excluded again below. The 30 healthy participants of the high-coverage sub-study (Cristiano et al., Table S5) were excluded because they were sequenced more deeply, are younger and are all female. Every remaining participant contributes one sequencing run, and the panel of normals shares no participant with the modeling set. Role, exclusion reason, outer fold and out-of-fold prediction of every record are given in Supplementary Table S1. BRCA, breast; PA, pancreatic; OV, ovarian; GA, gastric; CRC, colorectal; BD, bile duct; LU, lung cancer.

**Fig. S2:**
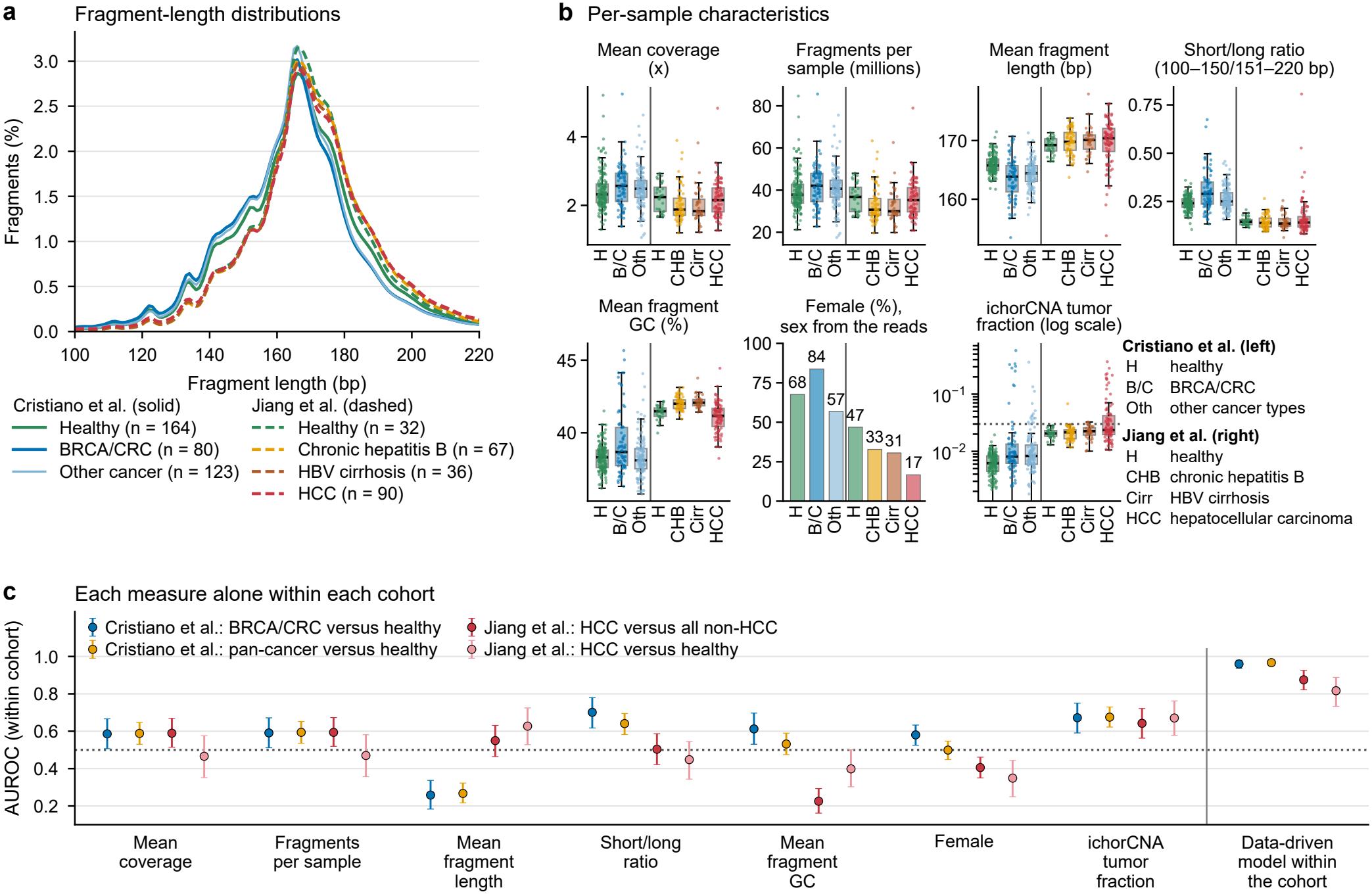
Comparison of the Cristiano et al. and Jiang et al. cohorts. a) Fragment-length distribution per cohort and clinical group, as the mean percentage of fragments at each length within the 100-220 bp window used for feature extraction (solid lines: Cristiano et al.; dashed lines: Jiang et al.). b) Per-sample characteristics per cohort and clinical group: mean genome-wide coverage, number of fragments, mean fragment length, short/long fragment ratio (100-150 bp / 151-220 bp), mean fragment GC content, percentage of female participants (sex inferred from the sex-chromosome reads), and ichorCNA tumor fraction (log scale; dotted line at 0.03), estimated in both cohorts with the same panel of normals from Cristiano et al. Boxes show the median and interquartile range; points are individual samples. c) AUROC of each characteristic alone for separating cancer from non-cancer participants within each cohort (BRCA/CRC versus healthy and pan-cancer versus healthy in the Cristiano et al. cohort; HCC versus all non-HCC participants and HCC versus healthy subjects in the Jiang et al. cohort; higher values indicating cancer, so that a value below 0.5 means that the characteristic is lower in the patients with cancer), next to the data-driven model trained and evaluated within the same cohort with nested cross-validation (pooled out-of-fold predictions). Error bars indicate 95% confidence intervals from 2,000 participant-level bootstrap resamples. H: healthy; B/C: BRCA/CRC; Oth: other cancer types (Cristiano et al.); CHB: chronic hepatitis B; Cirr: HBV-associated cirrhosis; HCC: hepatocellular carcinoma (Jiang et al.).

**Fig. S3:**
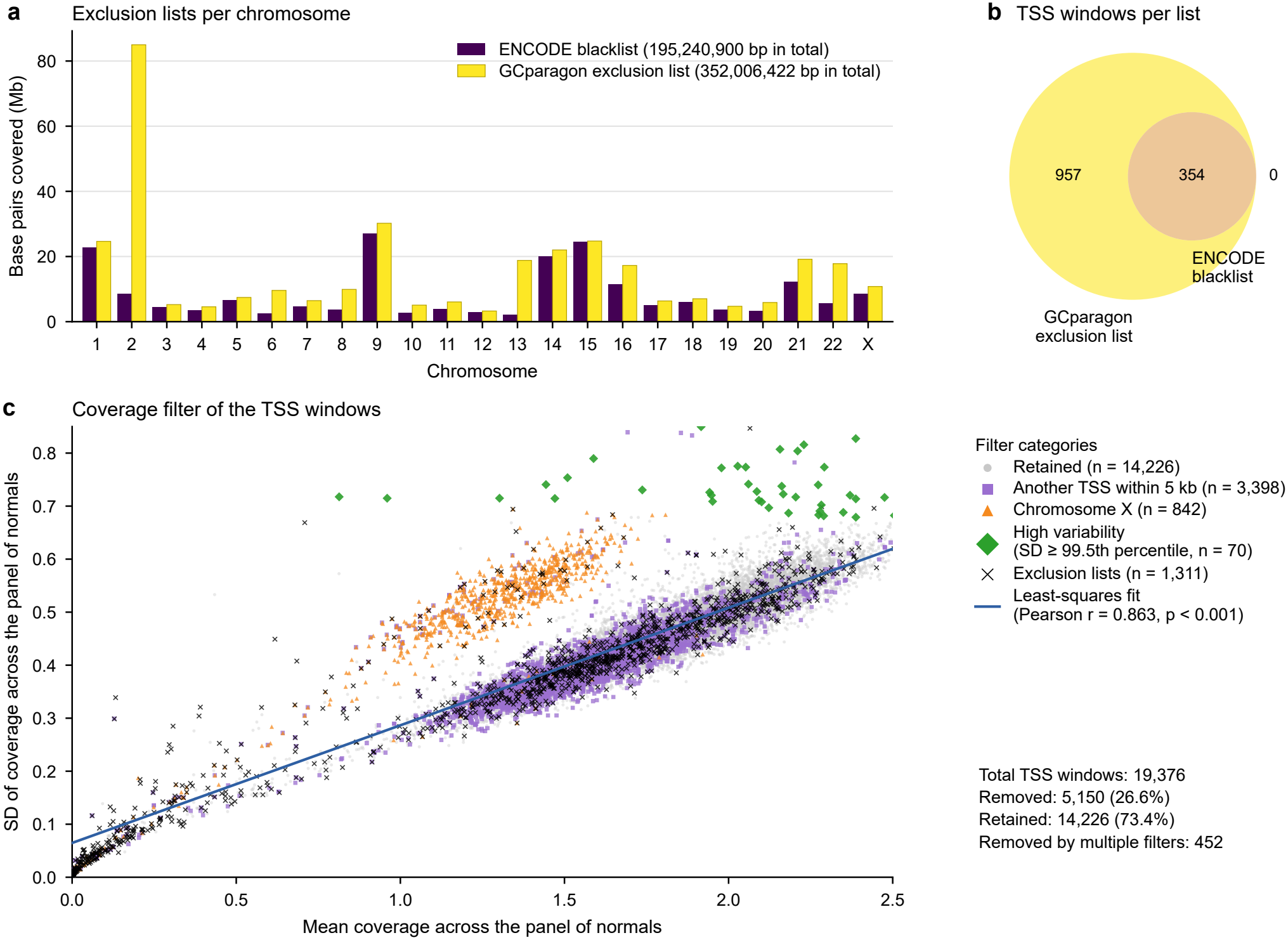
Quality control of the 10k bp TSS windows. **(a)** Base pairs per chromosome covered by the ENCODE blacklist and by the more conservative GCparagon exclusion list. **(b)** TSS windows that overlap each list; every window that overlaps the ENCODE blacklist also overlaps the GCparagon list, which flags 1,311 windows in total. **(c)** Mean and standard deviation of coverage of every TSS window across the 50 healthy donors of the panel of normals, with the least-squares fit (Pearson *r* = 0.863). Windows are removed if they overlap the exclusion lists, lie on chromosome X (coverage depends on sex), have another TSS within 5 kb, or have a standard deviation at or above the 99.5th percentile across the panel of normals (70 windows beyond the other filters). Of the 19,376 windows, 5,150 (26.6%) are removed, 452 of them by more than one filter, and 14,226 are retained. A few windows outside the plotted range are not shown; all of them are removed.

**Fig. S4:**
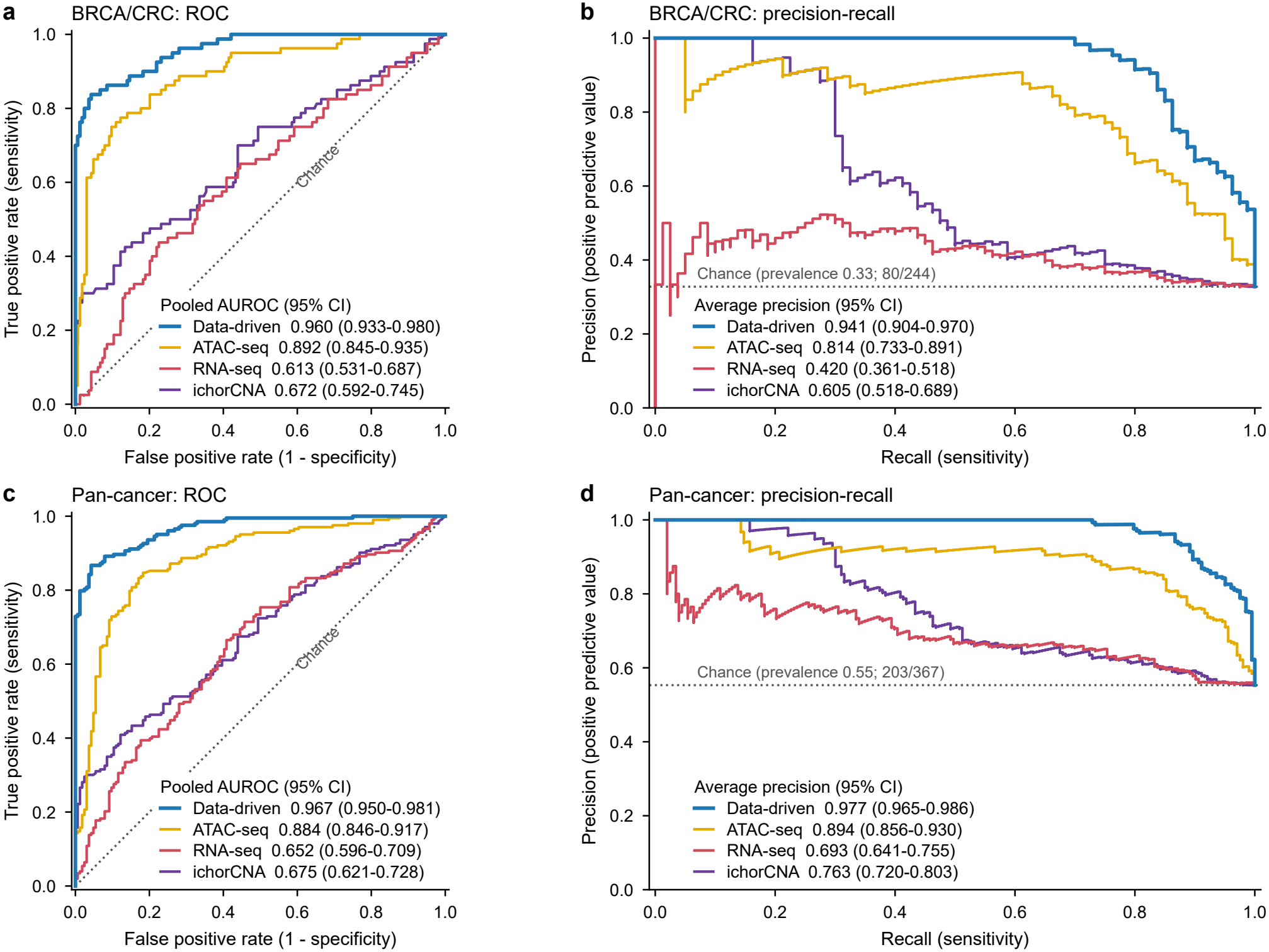
Pooled out-of-fold ROC and precision-recall curves. Every participant contributes the score from the one outer fold in which it was tested, and the four methods are scored on the same participants. a, b) BRCA/CRC versus healthy (80 cancers, 164 healthy donors). c, d) Pan-cancer versus healthy (203 cancers, 164 healthy donors). a, c) ROC curves; the legend gives the pooled AUROC with its participant-level bootstrap 95% confidence interval (2,000 resamples within each class); dotted line: chance. b, d) Precision-recall curves; the legend gives the average precision with its bootstrap 95% confidence interval; dotted line: the prevalence of cancer. These intervals give the uncertainty of the pooled estimate, whereas the shaded bands of the mean ROC curves in Figures 3a and 4a show the variability between the outer folds.

**Fig. S5:**
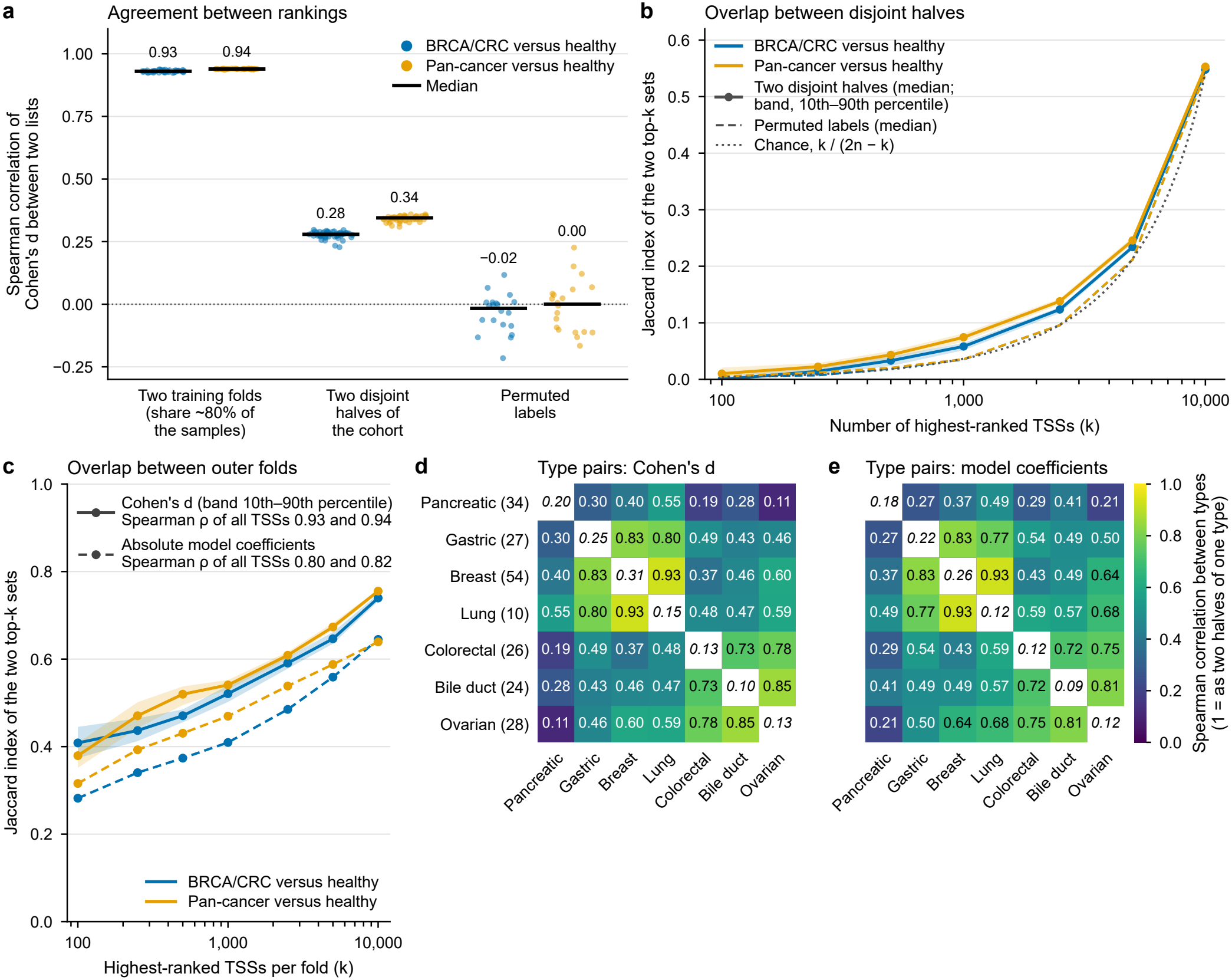
Detail of the TSS ranking: stability, model coefficients and pairs of cancer types. The 14,221 TSSs are ranked by the standardized difference in dip depth (GD) between patients with cancer and healthy donors (Cohen’s *d*); blue: BRCA/CRC versus healthy, orange: pan-cancer versus healthy. a) Spearman correlation of *d* between two rankings: those of the training samples of two outer folds, which have about 80% of the cohort in common (45 pairs); those of two halves of the cohort that share no samples (50 random splits); and the ranking made with permuted labels against the real one (20 permutations). Points: single comparisons; bars and numbers: medians. b) Overlap (Jaccard index) of the *k* highest-ranked TSSs of two rankings, for two disjoint halves of the cohort (solid: median; shaded: 10th to 90th percentile) and for permuted labels (dashed: median); dotted: the overlap expected by chance. c) Overlap of the *k* highest-ranked TSSs between the training samples of two outer folds (median over the 45 pairs of folds), for the ranking by *d* (solid; shaded: 10th to 90th percentile) and for the ranking by the absolute coefficients of the model fitted in each fold (dashed; a TSS that a fold did not select counts as zero). The legend gives the Spearman correlation over all TSSs, for BRCA/CRC versus healthy and pan-cancer versus healthy. d, e) Agreement of the ranking between pairs of cancer types, for *d* (d; 200 random splits) and for the coefficients of the pan-cancer model configuration (GD, all TSSs, elastic-net logistic regression) fitted on each half (e; 20 splits). In each split, the 164 healthy donors and the patients of every cancer type were divided into halves, so that two compared halves never share a sample. Diagonal (italic): Spearman correlation between the two halves of the same type. Off the diagonal: Spearman correlation between a half of one type and a half of another, divided by the geometric mean of the two within-type correlations, so that 1 means as similar as two halves of the same type. Number of patients per type in brackets.

**Fig. S6:**
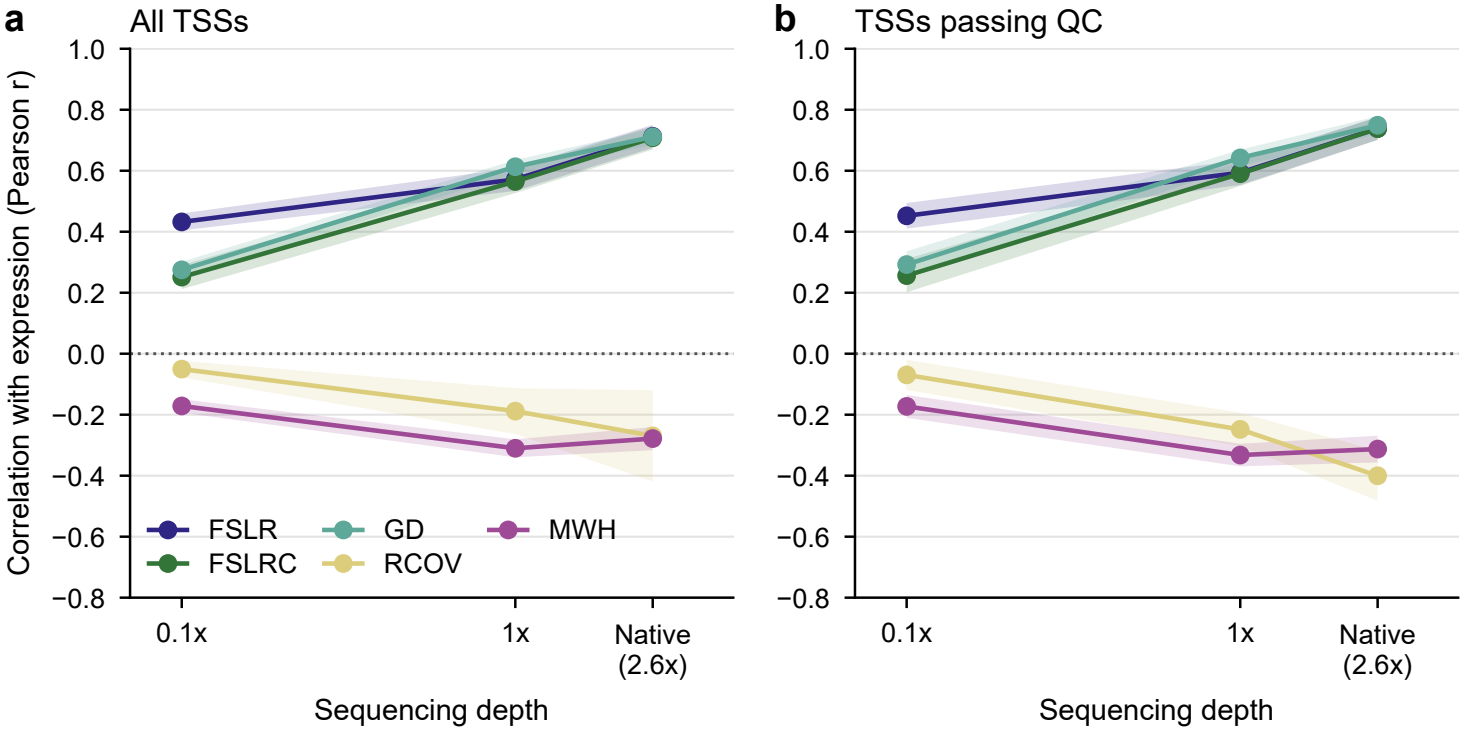
Correlation of the cfDNA features with gene expression as a function of sequencing depth. Pearson correlation of each feature with PBMC gene expression (as in Figure 1c) for the 50 samples of the panel of normals at their native depth (median 2.6x) and after downsampling each sample to 1x and 0.1x mean depth. Fragments were thinned at random with a fixed seed, so the same samples are compared at every depth. Lines show the mean over the 50 samples and shaded bands *±* 1 standard deviation. a) All TSSs. b) TSSsthat pass the quality filter. At 0.1x, more than 90% of the RCOV and MWH values are zero.

**Fig. S7:**
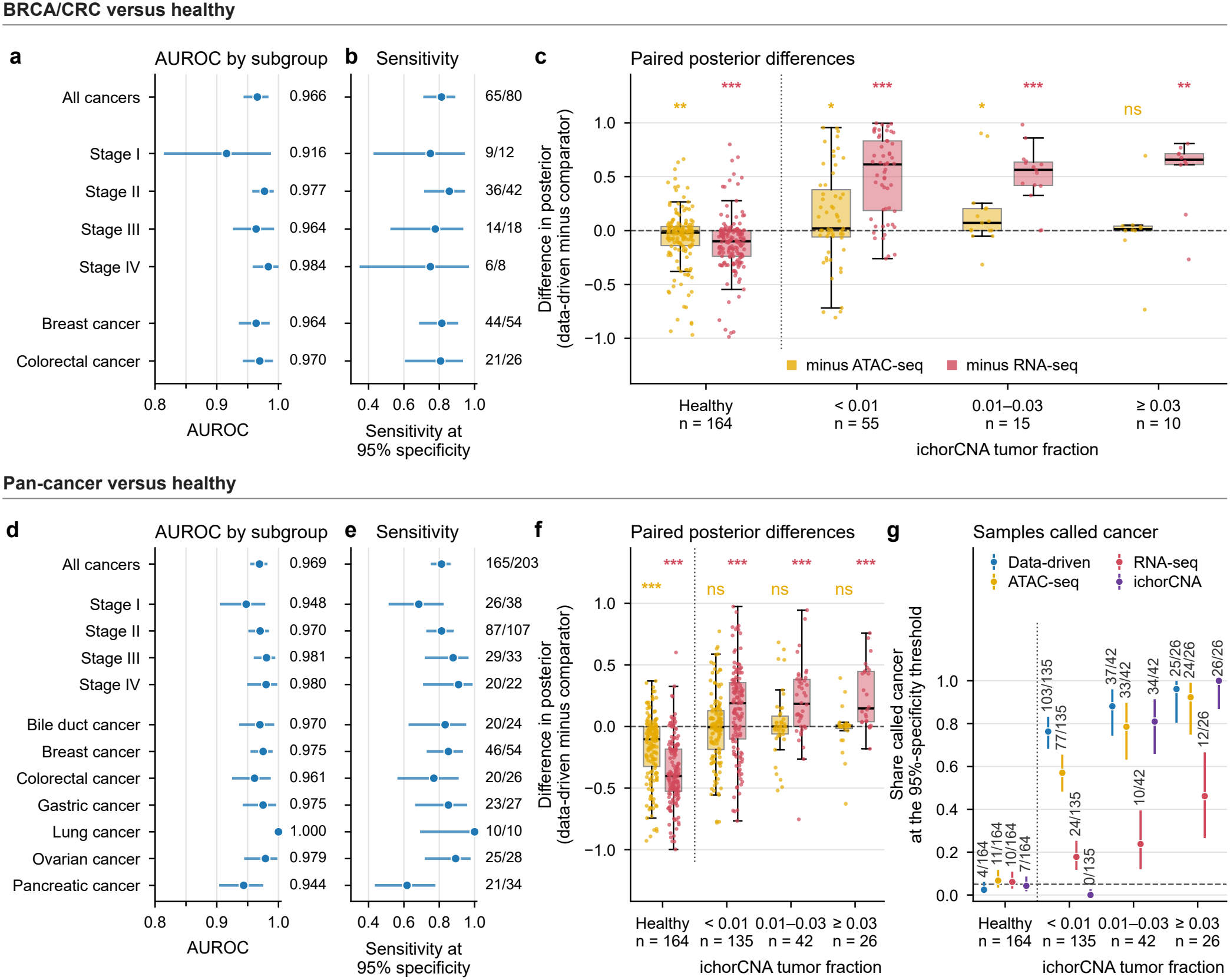
Performance by cancer type, stage and ichorCNA tumor fraction. Top row: BRCA/CRC versus healthy (80 cancers, 164 healthy donors); bottom row: pan-cancer versus healthy (203 cancers, 164 healthy donors). All values are out-of-fold predictions of the nested CV. Three patients of the pan-cancer task without a stage I to IV (two stage 0, one not staged) are in no stage group. a, d) AUROC of the data-driven model for all cancers, per stage and per cancer type, each against all healthy donors, with participant-level bootstrap 95% confidence intervals. b, e) Sensitivity for the same groups at the 95% specificity threshold derived from the training samples of each outer fold, with exact binomial 95% confidence intervals; numbers give detected / total. c, f) Paired differences in the out-of-fold posterior, data-driven model minus ATAC-seq or RNA-seq, for healthy donors and for cancers grouped by their ichorCNA tumor fraction (below 0.01, 0.01 to 0.03, 0.03 or above); positive values favor the data-driven model for cancers, negative values for healthy donors. Boxes show the median and interquartile range, points individual samples. One-sided Wilcoxon signed-rank tests in the direction favoring the data-driven model, not corrected for multiple comparisons (exploratory): ns, not significant; \**p <* 0.05; \*\**p <* 0.01; \*\*\**p <* 0.001. g) Share of samples called cancer by each method at its 95% specificity threshold derived from the training samples of each outer fold, with exact binomial 95% confidence intervals; numbers give called / total and the dashed line marks 5%. The same panel for BRCA/CRC versus healthy, with all cancers as an additional group, is Figure 3b. For ichorCNA the score is the tumor fraction itself, so it is not shown in c and f.

**Fig. S8:**
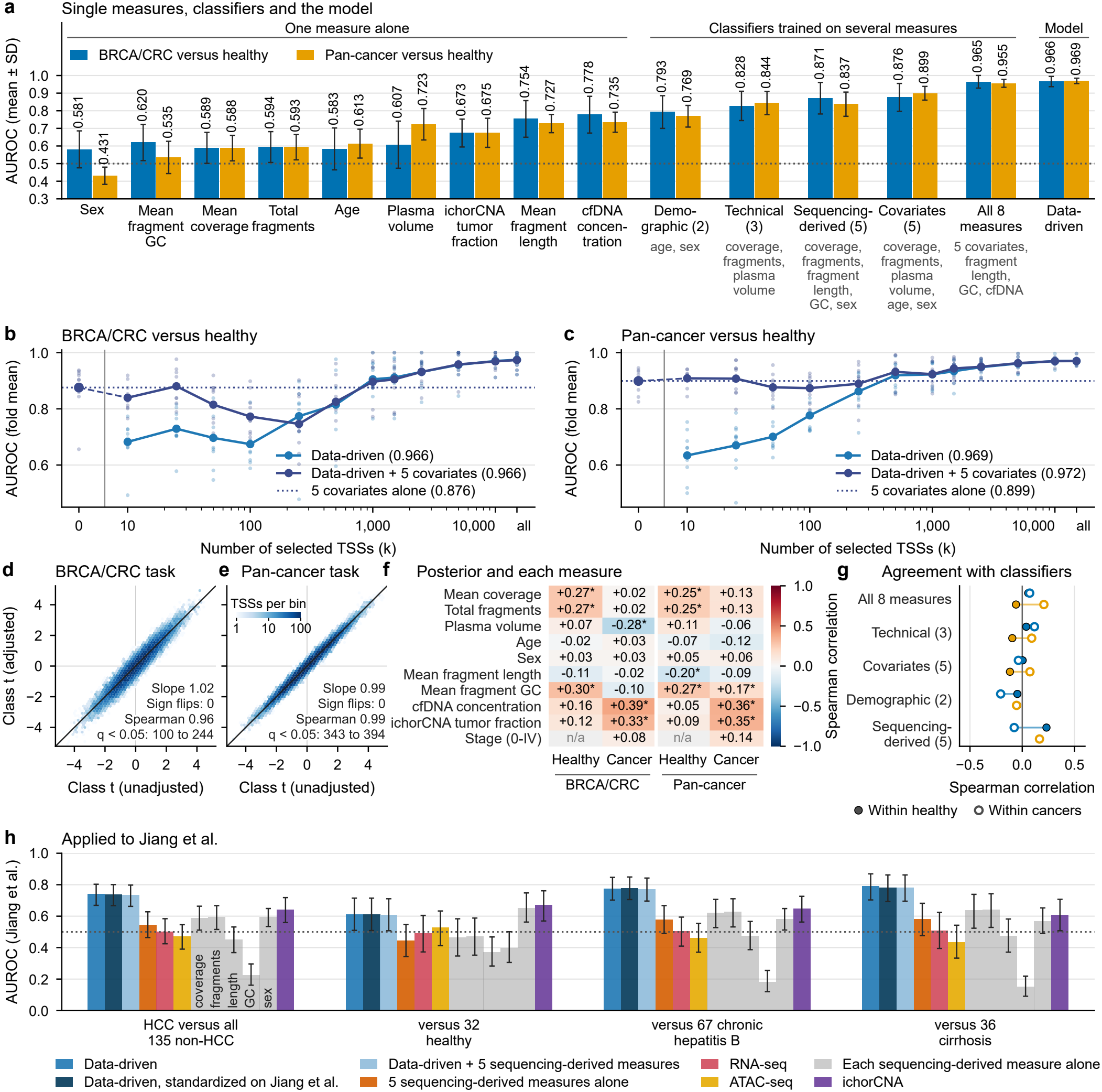
Technical and demographic covariates and the data-driven model, for BRCA/CRC versus healthy (blue; 80 cancers, 164 healthy donors) and pan-cancer versus healthy (orange; 203 cancers). a) AUROC (mean *±* SD over the ten outer folds) of each per-sample measure alone, of classifiers trained on the measures listed under each name (selected by the same nested procedure as the data-drivenmodel) and of the data-driven model; below 0.5, the direction set on the training folds reversed in the test folds. ichorCNA tumor fraction is shown for comparison only. b, c) AUROC against the number of selected TSSs (*k*) for the data-driven model alone and with the five covariates added as features (lines: mean; points: single folds; *k* = 0 and dotted line: covariates alone); legend: AUROC with *k* selected inside the folds. d, e) Class effect per TSS: *t*-statistic of class in a regression of the standardized GD on class alone (x-axis) and with coverage, plasma volume, age and sex added (y-axis), for the 14,221 TSSs; printed: their Spearman correlation and slope, the TSSs with BH *q <* 0.05 and sign flips among those. f) Spearman correlation of the out-of-fold score of the data-driven model with each measure, and with ichorCNA tumor fraction and stage, within healthy donors and within cancers (*: BH *q <* 0.05). g) Spearman correlation between the out-of-fold scores of the data-driven model and of each covariate classifier, within outer fold and class. h) Pan-cancer models trained on the Cristiano et al. cohort and applied once to the Jiang et al. dataset: AUROC for the 90 HCC patients against all 135 non-HCC participants and against each comparison group (participant-level bootstrap 95% confidence intervals), for the methods in the legend; gray: each sequencing-derived measure alone, named in the first group (same order in the others).

**Fig. S9:**
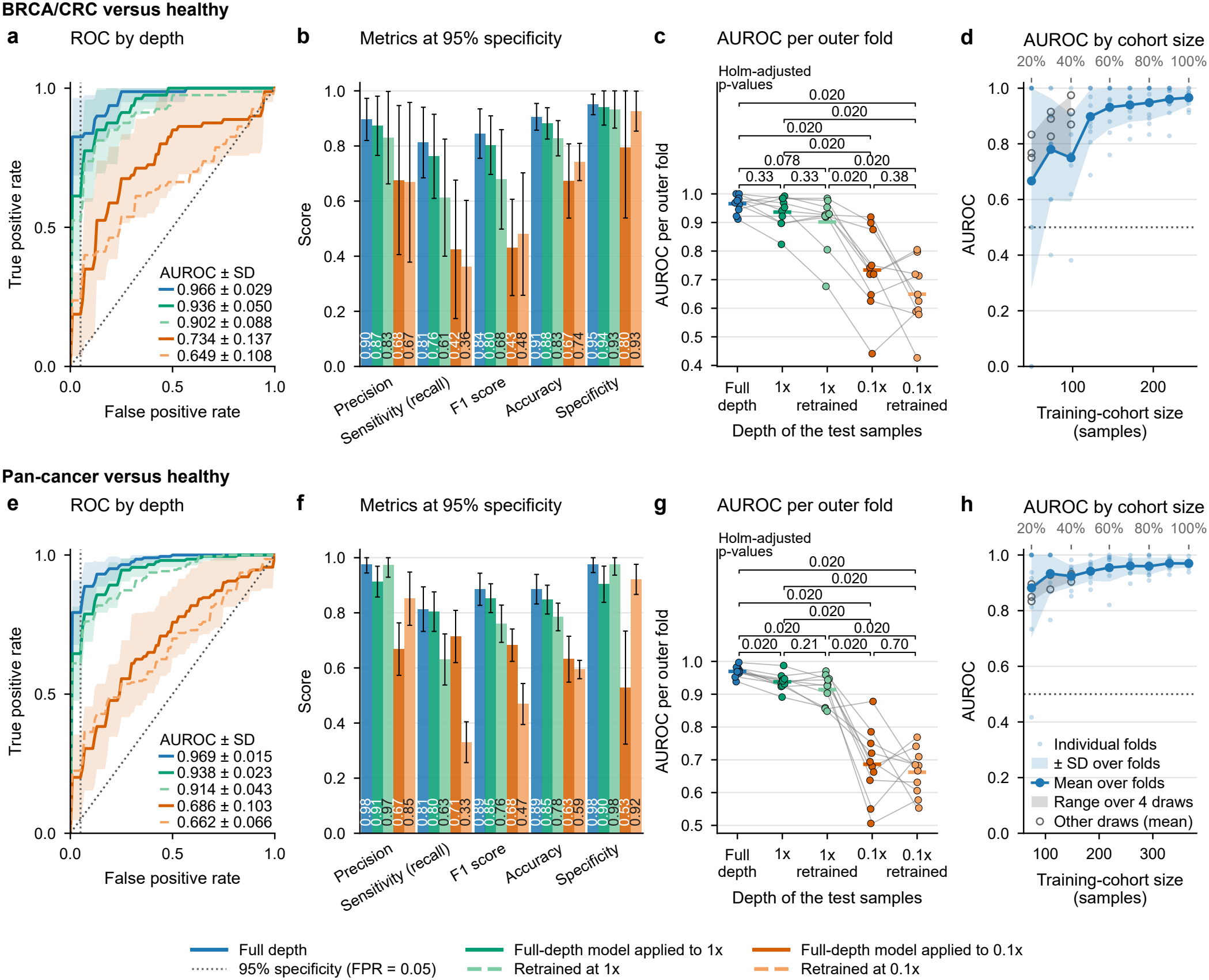
Robustness of the data-driven model to sequencing depth and training-cohort size. The top row shows BRCA/CRC versus healthy (a-d) and the bottom row shows pan-cancer versus healthy (e-h). a-c, e-g) All samples were downsampled to a uniform 1x or 0.1x coverage and compared with full depth (~2.48x). Solid lines and dark bars show the full-depth models applied unchanged to downsampled test folds. Dashed lines and pale bars show models retrained at the downsampled depth. a, e) Mean ROC curves over the 10 outer folds, with the shaded regions equaling one standard deviation. b, f) Classification metrics at the 95% specificity threshold on the cancer class. Error bars indicate one standard deviation over the 10 outer folds. The full-depth models keep their full-depth threshold, which does not transfer to lower depths, so their specificity drops. c, g) AUROC per outer fold, with lines connecting the same fold across depths and horizontal bars marking the mean. Brackets compare every two arms: two-sided Wilcoxon signed-rank test on the paired fold AUROCs, Holm-adjusted over the ten pairs (with ten folds, the smallest attainable adjusted p-value is 0.020). d, h) AUROC against training-cohort size. The complete nested cross-validation (10 outer folds) was repeated on stratified subsets of 20% to 100% of the cohort, each subset contained in the next larger one. Shown are the mean (solid line) and standard deviation (shaded) over the outer folds and the individual folds (light points), against the number of samples, with the percentage of the cohort on the top axis. The subsets of 20% to 40% were drawn three more times (open circles: mean AUROC of each draw); the gray band spans the mean AUROC over the four draws.

**Fig. S10:**
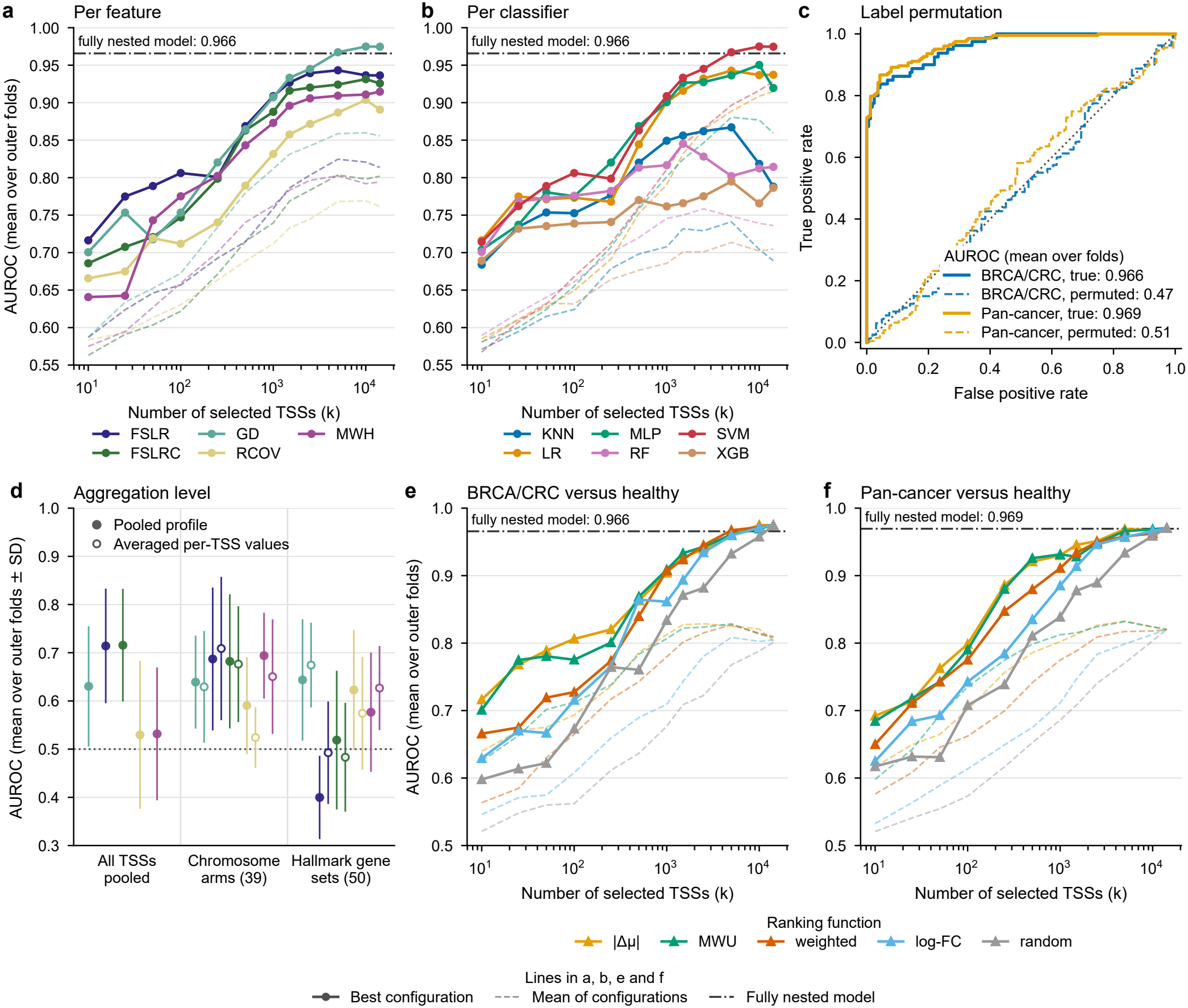
Analysis of the data-driven model for BRCA/CRC versus healthy, and the ranking functions in both tasks. a, b, d) BRCA/CRC versus healthy (80 cancers, 164 healthy donors). In a, b, e and f, every configuration of the data-driven model (feature, ranking function, number of selected TSSs *k* and classifier, with the hyperparameters tuned in the inner loop) was scored on the outer test folds; shown is the AUROC (mean over the 10 outer folds) against *k*. Solid lines: the best configuration with a given feature (a), classifier (b) or ranking function (e, f) at that *k*; dashed lines: the mean over these configurations. These curves are descriptive; the dash-dotted line marks the AUROC of the fully nested model, in which the configuration is chosen inside each outer fold (0.966 for BRCA/CRC versus healthy, 0.969 for pan-cancer versus healthy). a) Per fragmentomic feature. b) Per classifier (KNN: K-Nearest Neighbors; LR: Logistic Regression; MLP: Multi-Layer Perceptron; RF: Random Forest; SVM: Support Vector Machine; XGB: XGBoost). c) Label permutation in both tasks: pooled out-of-fold ROC curves of the fully nested procedure with the true labels (solid) and with the diagnosis labels permuted once before the outer folds were drawn (dashed), for BRCA/CRC versus healthy (blue) and pan-cancer versus healthy (orange; 203 cancers). The legend gives the AUROC as the mean over the 10 outer folds (0.966 and 0.969 with the true labels, 0.47 and 0.51 with permuted labels); dotted line: chance. d) AUROC by level of aggregation: each fragmentomic feature computed on all TSSs pooled (one value per sample), on the 39 autosomal chromosome arms (57 to 845 TSSs each) and on the 50 MSigDB Hallmark gene sets (29 to 177 TSSs each), from the pooled coverage profiles or fragment counts (filled) or, for the arms and gene sets, as the average of the per-TSS values (open). The features of the arms and of the gene sets went through the same nested procedure on the same outer folds; the single value per sample was used as the score itself, with its direction (higher or lower in cancer) set on the training samples of each outerfold. Points show the mean and lines the standard deviation over the 10 outer folds; the dotted line marks an AUROC of 0.5. e, f) Per ranking function, for BRCA/CRC versus healthy (e) and pan-cancer versus healthy (f; 203 cancers, 164 healthy donors). |Δ*μ*|: absolutedifference between group means; MWU: p-value of the Mann-Whitney U test; weighted: weighted log-ratio; log-ratio: absolute log_2_ ratio;random (gray): random selection of *k* TSSs, a comparator and not a ranking function. With all 14,221 TSSs nothing is selected, so the five curves meet.

**Fig. S11:**
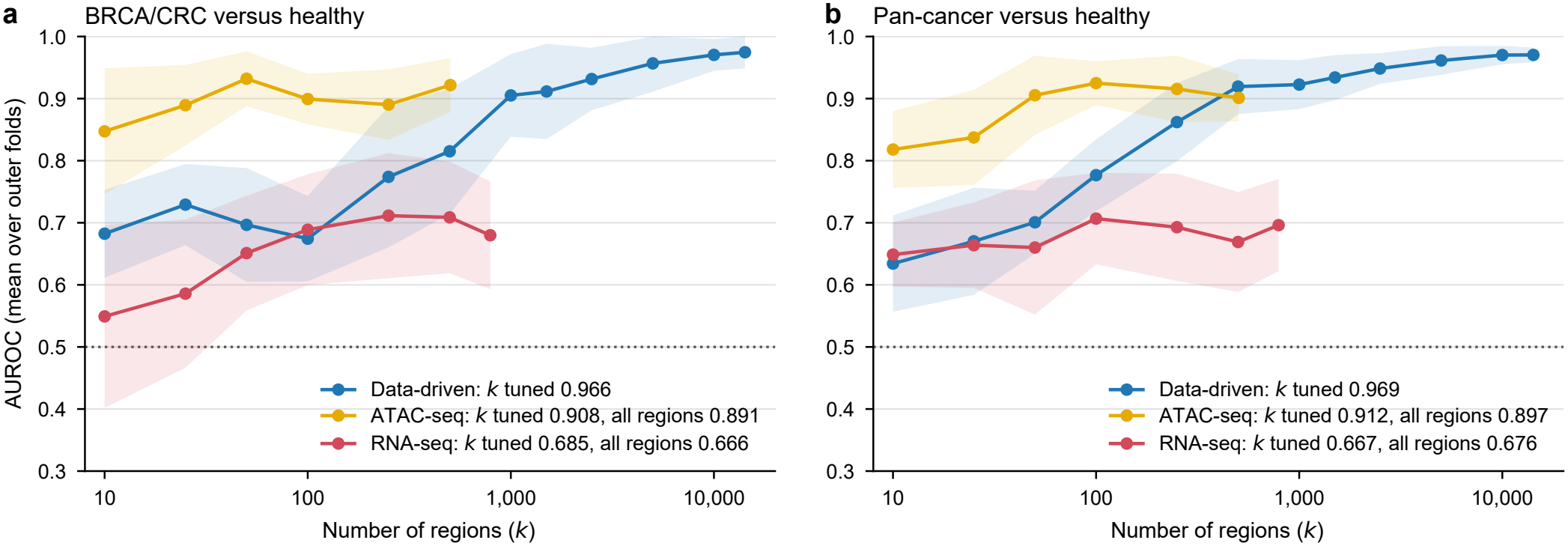
Region-number sweep for every method, BRCA/CRC versus healthy (a) and pan-cancer versus healthy (b). AUROC against the number of regions *k* for the data-driven model (TSSs), the ATAC-seq baseline (TFBS sets) and the RNA-seq baseline. Each point isa complete, fully nested run at that fixed *k* (mean over the ten outer folds; shaded: *±* SD); the last point of each curve uses all regions of that method. This point is its own run of the nested procedure and can differ from the baseline of Figures 3 and 4 (“all regions” in the legend), which was run separately. Because these curves are read off the outer test folds, their peaks are optimistic; the valid estimate is the AUROC with *k* chosen in the inner loop (“*k* tuned” in the legend), shown next to the AUROC with all regions.

**Fig. S12:**
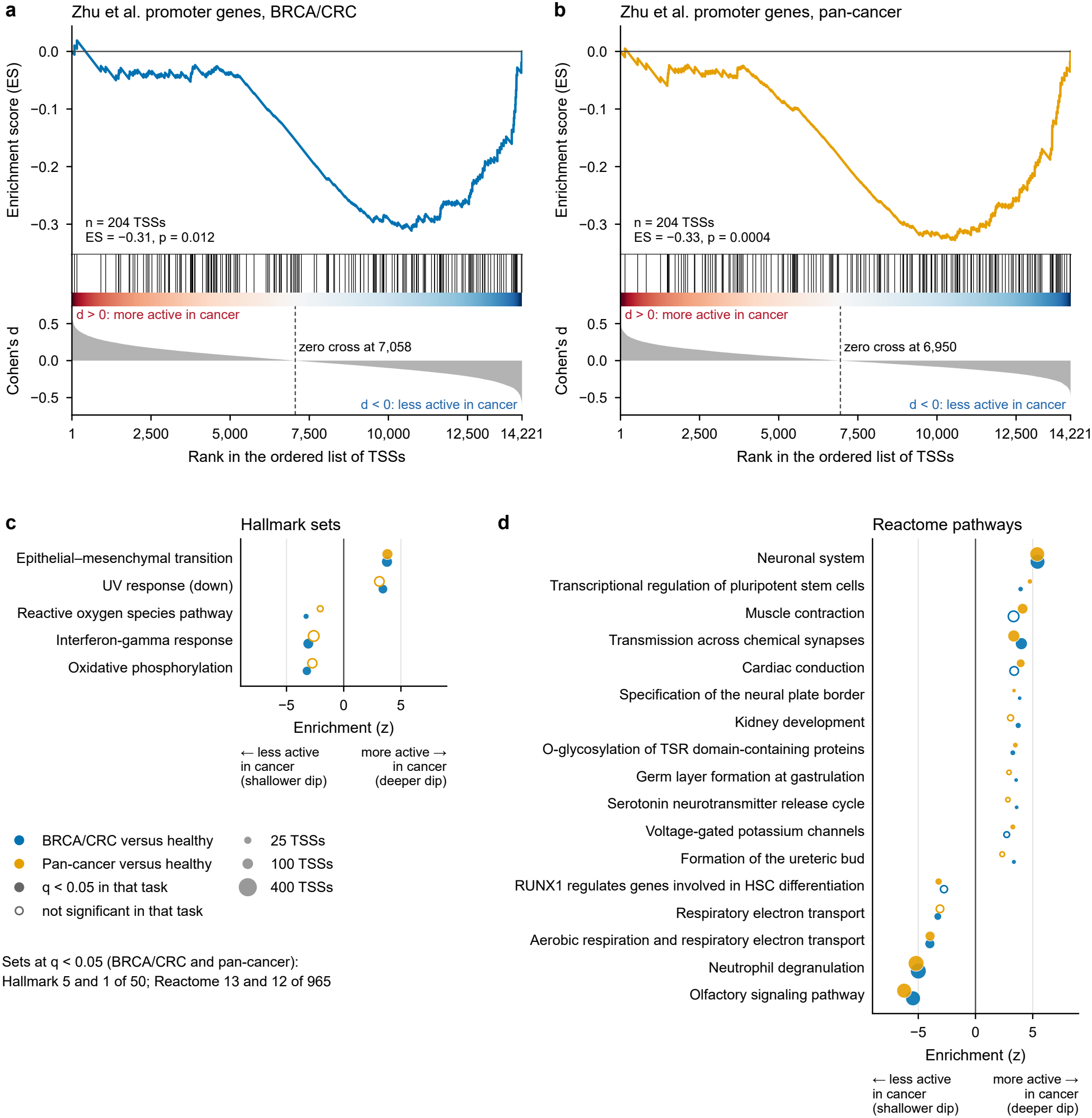
Gene sets along the ranked list of TSSs. The 14,221 TSSs were ranked by the standardized difference in dip depth between patients with cancer and healthy donors (Cohen’s *d*; positive: deeper dip in cancer, a more active promoter), for BRCA/CRC versus healthy (blue; 80 cancers, 164 healthy donors) and pan-cancer versus healthy (orange; 203 cancers). Each gene set was compared with 10,000 random sets of TSSs of the same size and the same composition over dip depth, coverage and GC content in healthy plasma. a, b) Enrichment of the 204 TSSs of the blood-specific promoter genes of Zhu et al. [24], for BRCA/CRC versus healthy (a) and pan-cancer versus healthy (b). Top: running enrichment score of a modified GSEA, with the TSSs ranked from the most active in cancer (left, *d >* 0) to the least active (right, *d <* 0); the most negative point of the curve is the enrichment score (ES). Middle: positions of these TSSs in the list (black lines) above the color scale of *d*. Bottom: *d* of every TSS along the list; the dashed line marks where *d* changes sign. *p*: two-sided empirical *p* of the ES among the random sets. c, d) MSigDB Hallmark sets (c; 50 tested) and Reactome pathways (d; 965 tested). Enrichment (*z*) is the position of the set’s TSSs in the ranked list relative to the random sets, in units of their standard deviation; *q*: two-sided empirical *p*, Benjamini-Hochberg corrected within a collection and task. All sets with *q <* 0.05 in at least one task are shown (filled: *q <* 0.05 in that task; open: not); dot area: number of TSSs in the set. The cell-type signatures tested in the same way are summarized in Figure 6b.

